# Hearing hippocampal ripples: Event-faithful, reproducible sonification of multineuronal spiking with hierarchy-guided orchestration

**DOI:** 10.64898/2026.09.05.749637

**Authors:** Yuji Ikegaya

**Author notes:** To whom correspondence should be addressed: Yuji Ikegaya, Ph.D. Laboratory of Chemical Pharmacology, Graduate School of Pharmaceutical Sciences, The University of Tokyo, 7-3-1 Hongo, Bunkyo-ku, Tokyo 113-0033, Japan.

## Abstract

Among the human senses, hearing is distinguished by its high frequency resolution and sensitivity to spatiotemporal patterns. Accordingly, sonification—the rendering of neural activity as sound for direct perception—has a long history in neuroscience. Although sonification can facilitate exploration of the temporal structure of population activity, dense spike trains are often difficult to track by ear. Here, I present HERO (Hierarchy-guided Event-faithful Reproducible Orchestration), a pipeline that assigns stable musical voices to recorded units while preserving spike onsets in a MIDI event representation. Each spike within a selected interval produces a single Note On message whose timing is determined by a global temporal dilation followed by bounded clock rounding. A separate analysis pathway uses multiscale correlations and block-bootstrap co-association to organize units into a hierarchy. This hierarchy guides instrument assignment, whereas each unit is assigned a fixed key, tuning offset, velocity, and stereo position. A derived SoundFont stores tuning and pan parameters independently for units that share a MIDI channel. Rate-dependent velocity and channel-volume settings are intended to reduce the dominance of frequently firing units. An online audio example illustrating the audibility of individual activity during large-scale firing is available at: https://www.youtube.com/watch?v=zOYGXcp727A. HERO provides a reproducible auditory representation of population activity that may help direct attention toward features warranting subsequent quantitative analysis.

## Introduction

Listening has long been part of electrophysiology. Adrian and Matthews (1934) described auditory monitoring of cortical electrical activity, and audio monitors remain useful during extracellular unit recording. Hearing is sensitive to temporal pattern, and differences in pitch, timbre, and spatial position can help listeners organize concurrent sounds into perceptual streams (Bregman, 1990). Sonification can therefore complement a raster plot by presenting activity as an unfolding sequence.

Neural sonification has served both artistic and scientific purposes (Lutters and Koehler, 2016). In Music for Solo Performer (1965), Alvin Lucier used amplified alpha activity to excite percussion instruments (Straebel and Thoben, 2014). Subsequent biofeedback compositions and brain–computer music interfaces also used EEG features to control sound. Such mappings can convey changes in neural activity without representing individual spikes or preserving every detected event. Their suitability depends on the signal and the purpose of the display.

Event-based approaches have a separate history. Aertsen and Erb (1987) assigned distinct tones to simultaneously recorded units in the neurophone. Hermann et al. (2002) developed sonification methods for EEG inspection, and Baier, Hermann and Stephani (2007) introduced real-time sonification of detected EEG rhythms; Väljamäe et al. (2013) reviewed this broader literature. The Spikiss project assigned neuronal activity to different instruments, including separate instruments for excitatory and inhibitory populations (Destexhe and Foubert, 2018). Destexhe and Foubert (2022) described a mapping in which each detected EEG motif triggered a sound at its onset, with envelope parameters derived from motif morphology.

For multineuronal spike trains, the challenge is to represent many concurrent point processes whose firing rates may differ greatly. Parameter-mapping sonification provides a general framework for choosing auditory attributes (Kramer et al., 1999; Hermann, Hunt and Neuhoff, 2011; Dubus and Bresin, 2013). Pitch, timbre, and spatial cues can support stream segregation, but they do not ensure that a listener can distinguish every source in a dense mixture (Bregman, 1990). A useful method must specify both what information it preserves and how the remaining sound-design choices are made.

Snapping onsets to a musical grid or omitting spikes changes the event sequence. Adjusting a fixed voice or gain does not necessarily change that sequence, although it changes the prominence of individual events. This distinction motivates the present design: onset timing is constrained separately from the orchestration used to present it.

HERO implements this separation with two pathways (Figure 1). The event pathway retains the input timestamps and converts spikes within a selected interval to MIDI Note On messages using a reported clock and playback rate. The analysis pathway bins and filters a copy of the raster to estimate temporal similarity and assign voices. A hierarchy derived from multiscale correlations and bootstrap co-association guides instrument assignment. Each unit then receives a fixed key, tuning offset, velocity, and pan. Tuning and pan are stored in a derived SoundFont, allowing units on the same MIDI channel to retain separate settings. The source data, mapping, configuration, and output digests provide a record against which event preservation can be checked.

**Figure 1.**
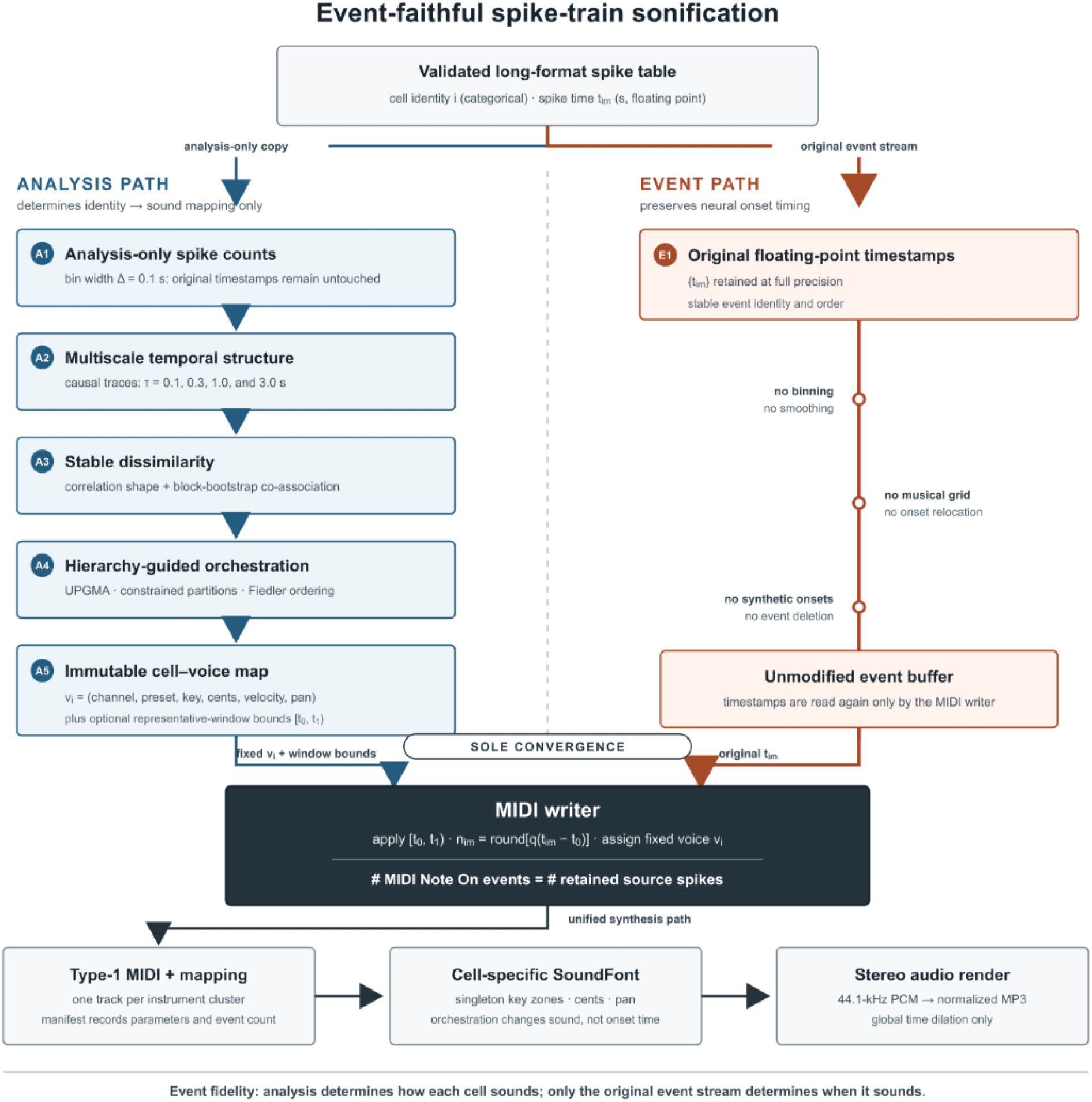
Separate analysis and event pathways. The analysis pathway bins a copy of the raster, computes multiscale correlations and bootstrap co-association, and constructs a hierarchy and fixed voice mapping. The event pathway retains the original timestamps for window selection and MIDI construction. Each selected spike produces one Note On at its clock-rounded time. The schematic describes data flow, not the number of memory accesses made by the code. SoundFont synthesis and audio encoding follow MIDI construction, so preservation of the event list does not establish acoustic onset fidelity. The schematic was generated with gpt-image-2 and edited manually.

The example uses hippocampal activity recorded from one rat in its home cage. Sharp-wave ripples recruit coordinated populations of pyramidal cells and interneurons (Csicsvari et al., 1999; Buzsáki, 2015), and activity during some such events reflects sequences expressed during behavior (Lee and Wilson, 2002; Foster and Wilson, 2006). A source interval of 50–100 ms occupies 1–2 s at the twentyfold dilation used here. Hippocampal pyramidal cells also generate short sequences of closely spaced spikes (Ranck, 1973; Harris et al., 2001), which motivate a fixed voice for each unit. However, the present analysis uses spike times alone: its population-burst labels do not establish ripple identity or replay. The listening observations are descriptive and identify candidate features for subsequent quantitative and perceptual tests.

The following sections specify the mapping, clock conversion, reference implementation, and verification procedures. Reproducibility is considered separately for the MIDI event representation, the voice mapping, and the synthesized audio.

## Methods

### Methodological overview and invariants

HERO converts a spike raster into a polyphonic, microtonal, spatialized rendering. The input contains N recorded unit identifiers and M spikes over an observation interval of duration T. The reference implementation requires at least two units with suitable temporal variation for clustering; N, T, and M are obtained from the input. The term cell in the code and figures denotes an input unit identifier and does not independently establish that each identifier corresponds to a distinct biological neuron. The one-port implementation uses at most 16 instrument clusters. Multiple ports or synchronized files are possible extensions but are not implemented in the supplied code.

Event fidelity is defined at the MIDI level. Within the selected source interval, each input spike must correspond to one Note On message. Onsets are neither moved to a musical grid nor selectively omitted; they undergo only the clock conversion in Eqs. 23–25. Statistical analysis determines a fixed mapping from unit identity to timbre, key, tuning offset, velocity, and pan. This preserves the encoded event sequence within the stated clock precision, but it does not guarantee that every event can be heard separately.

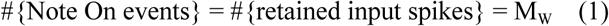

Each unit i is assigned a fixed voice tuple v_i_ = (port, channel, preset, key, cents, velocity, pan), used for all of its spikes. The reference implementation uses one port. Unit identifiers are categorical labels rather than numerical pitch values, although their initial ordering can affect deterministic tie-breaking.

### Input data model and validation

The input is a long-format CSV file with one row per spike and two required fields: a unit identifier and a timestamp. Let t_im_, m = 1, …, M_i_, denote the ordered timestamps of unit i. Acquisition start t_start_ and exclusive stop t_stop_ should be supplied because silent recording margins affect firing-rate estimates.

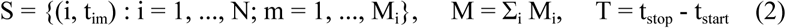

The loader rejects missing identifiers, missing or non-finite timestamps, negative timestamps, exact duplicate (unit, time) pairs, and inputs with fewer than two units. Recording bounds must be finite and satisfy t_start_ ≤ t_im_ < t_stop_ for every spike. Timestamps in a column whose name ends in _ms are divided by 1,000; other time columns are interpreted as seconds. The command line accepts custom column names and recording bounds. Rows are sorted stably by time and unit label, and unit labels are ordered lexicographically for matrix construction. If bounds are omitted, the code uses t_start_ = min(S.t) and t_stop_ = nextafter(max(S.t), +∞). This retains the last spike but excludes unrecorded silent margins from the rate denominator. Within-unit timestamps are sorted before interspike intervals are calculated.

### Separation of the analysis and event pathways

The analysis pathway bins and filters a copy of the raster to estimate similarity and construct the voice mapping. The event pathway retains the original floating-point timestamps and uses them for source-window selection and MIDI construction. Analysis bins therefore do not determine MIDI onset times.

1. Analysis pathway: spike raster → binned counts → multiscale traces → dissimilarity and hierarchy → fixed unit-to-voice mapping.
2. Event pathway: original timestamps → source-window selection → clock conversion → MIDI Note On messages.
3. Synthesis pathway: mapping and MIDI → unit-specific SoundFont zones → stereo PCM → normalized MP3.

### Analysis-only binning and multiscale spike-density traces

For clustering only, the recording was partitioned into *B* bins of width Δ seconds (default Δ = 0.1 s). The integer count for cell *i* in bin *n* was

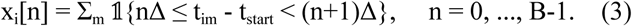

To capture coordinated structure at more than one temporal scale, each count train was passed through four causal exponential filters with time constants τ*_l_* ∈ {0.1, 0.3, 1.0, 3.0} s. With α*_l_* = exp(-Δ/τ*_l_*), the recursion was initialized at zero and evaluated as

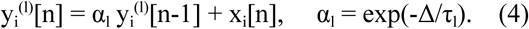

Each filtered trace was standardized across time with the population mean and population standard deviation. A cell with zero temporal variance after filtering is undefined for correlation clustering and therefore triggers an explicit error.

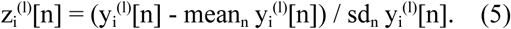

### Multiscale correlation dissimilarity

For each scale *l*, the pairwise correlation was the normalized inner product of the standardized traces. Because population standardization was used, the diagonal is one up to floating-point error and was set exactly to one.

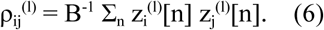

The scale-specific correlations were combined with non-negative weights *w* = (0.35, 0.30, 0.20, 0.15), which sum to one. The resulting chord distance lies in [0,2] because the weighted similarity lies in [-1,1].

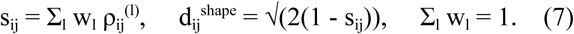

### Block-bootstrap stability and consensus dissimilarity

A moving-block resampling procedure was used to estimate co-association. For each of R replicates (default 12), Q blocks (default 10) of duration L_b_ = 10 s were sampled with replacement from valid starting bins of the already filtered, standardized traces. The same sampled bins were used for all units and scales. Concatenated trace blocks were re-centered and re-standardized, and correlations and d_shape_ were recomputed before a K-cluster UPGMA cut. The traces were not filtered again at block boundaries. If a unit had zero variance in a resample, its centered trace was left at zero by setting the divisor to one.

Thus each default replicate uses 100 s of sampled trace, rather than a full-length resample of the 3,661-s recording. C_ij_ denotes the fraction of replicates in which units i and j were assigned to the same cluster:

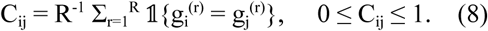

Let *m*_d_ be the median of the strictly positive upper-triangular values of *d*_shape_. The final symmetric dissimilarity combined normalized temporal shape and failure to co-cluster, with λ = 0.80.

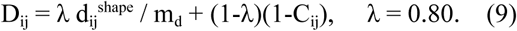

*D* is used as a dissimilarity; the method does not require the consensus term to satisfy all metric axioms. The diagonal is set to zero and numerical symmetry is enforced by averaging *D* with its transpose.

### UPGMA hierarchy and constrained partitioning

Agglomerative clustering used the unweighted pair-group method with arithmetic averages (UPGMA; Sokal and Michener, 1958). At each step, the two active clusters with the smallest dissimilarity were merged. For another active cluster C, the update was

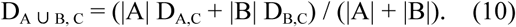

Ties were resolved deterministically by row-major node order. The binary tree was first cut into *G* broad superclusters (default *G* = 5) by dynamic programming constrained to tree nodes. The cost assigned to a candidate group *A* was the within-group squared-dissimilarity sum normalized by group size,

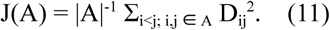

Within each supercluster, a tree-consistent leaf order was partitioned into contiguous instrument clusters. These finer clusters need not themselves be dendrogram clades. A second dynamic program allocated k_g_ clusters to supercluster g so that Σg k_g_ = K (default 16), every unit was assigned once, no cluster exceeded m_eff_, and the total cost J was minimized. The search began at m_eff_ = max(m_user_, ⌈N/K⌉), with a default soft capacity m_user_ = 15, and increased m_eff_ until a feasible allocation was found. The realized capacity was recorded in the manifest. Both dynamic programs retain the first encountered solution when costs tie exactly.

### Spectral orientation and cell ordering

A graph-spectral coordinate was used to orient otherwise reversible dendrogram branches. The Gaussian affinity matrix was computed from D using σ equal to the median positive pairwise dissimilarity; self-affinities were removed. The combinatorial graph Laplacian was then diagonalized.

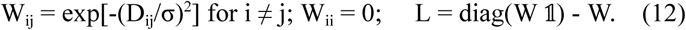

The Fiedler coordinate f was the eigenvector associated with the second-smallest eigenvalue of L (Fiedler, 1973). Its sign was reversed when its inner product with centered burst fractions was negative. At each dendrogram bifurcation, the child with the smaller mean coordinate was placed first; ties retained the existing child order. This ordering controlled key rank and within-cluster pan, never event time. The sign rule does not uniquely orient a vector orthogonal to the burst descriptor, nor does it resolve a repeated second eigenvalue. Consequently, identical spectral ordering across numerical environments is not guaranteed; the realized mapping must be retained.

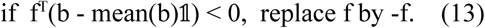

### Cell- and group-level descriptors

Descriptors were calculated over the full recording. Firing rate was r_i_ = M_i_/T. For interspike intervals I_im_ = ti,m+1 − t_im_, b_i_ was the fraction shorter than 50 ms, and local irregularity was measured by CV2 (Holt et al., 1996). The code calls b_i_ burst_fraction, but it is an operational short-ISI fraction, not a detector of canonical complex-spike bursts (Ranck, 1973; Harris et al., 2001). In the reported dataset, 732,652 of 1,501,745 within-unit ISIs (48.79%) met this criterion. The comparison is performed on floating-point timestamps in seconds; comparing the original millisecond values gives 732,586 intervals because of numerical classification at the 50-ms boundary. The seconds-valued comparison is used throughout the reference analysis.

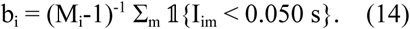

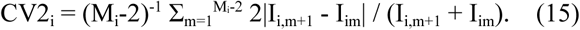

When too few intervals were available, the corresponding descriptor was set to zero. Group descriptors used the arithmetic means of unit firing rate, short-ISI fraction, and CV2, together with mean off-diagonal correlations at the shortest and longest scales and mean bootstrap co-association. Single-unit groups were assigned correlation and stability values of one by convention. These singleton values are bookkeeping choices, not empirical estimates of coherence.

### Mapping superclusters to timbral families

With G = 5, superclusters were assigned to five timbral families by a greedy rule. Scores were standardized once across all superclusters before assignment. The first four rules were applied in the order below, each time excluding previously assigned groups; the remaining group received the mallet label. If fewer than five groups were available, assignment stopped when one remained, and that group received the mallet label. With more than five groups, all groups remaining after the first four rules received the mallet label. Exact score ties retained the first group in the stored order.

4. Percussive: maximize *z*(log_10_ mean rate) + 0.7 *z*(short-scale coherence).

5. Sparse wind: minimize log_10_ mean rate among remaining superclusters.

6. Resonant metal: maximize *z*(long-scale coherence - short-scale coherence).

7. Plucked string: maximize *z*(mean burst fraction) + 0.4 *z*(mean CV2).

8. Mallet: assign the remaining supercluster.

Instrument selection began with the family assigned to each supercluster. Within a family, instrument clusters were ranked by short-scale coherence for percussion, short-ISI fraction for plucked instruments, long-scale coherence for resonant metal, and mean firing rate for mallets and sparse winds. Ties were broken by node index. Instruments were assigned from the ordered pools below without reuse across the ensemble. When a pool was exhausted, the first unused instrument in the complete palette was used. GM Percussion normally went to the first percussive cluster in this ranking; a fallback assigned it to the highest-rate percussive cluster only if no percussion kit had yet been selected. The mapping records the family of the actual instrument, which can differ from the parent supercluster family when a fallback instrument is used.

| Family | Ordered instrument pool | Within-family ranking |
| --- | --- | --- |
| Percussive | GM Percussion; Taiko Drum; Woodblock; Melodic Tom; Agogo; Xylophone | Short-scale coherence |
| Plucked | Koto; Shamisen; Sitar; Dulcimer; Kalimba | Burst fraction |
| Resonant metal | Vibraphone; Celesta; Tubular Bells; Tinkle Bell; Glockenspiel; Agogo | Long coherence |
| Mallet | Marimba; Xylophone; Kalimba; Dulcimer; Woodblock | Mean rate |
| Sparse wind | Shakuhachi; Shanai; Ocarina; Pan Flute; Koto | Mean rate |

### Optional selection of a representative source window

The full recording can be rendered by setting --window-s to at least T; Config.window_s = None also selects the full interval when using the Python API. For an excerpt of W seconds, candidate starts are sampled at stride h (defaults W = 30 s and h = 5 s). Let n_i_(w) and n_k_(w) be unit-wise and cluster-wise counts, y_s_(w) be one-second population counts, M_w_ be the total, and M_med_ be the median total over candidates. The active fraction and normalized entropies are

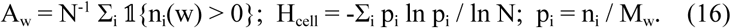

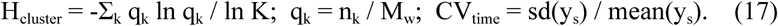

Zero-count categories were omitted from entropy sums. The selected window maximized

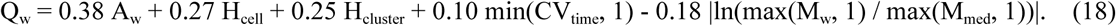

The score favors broad participation and temporal contrast while penalizing excerpts with atypical spike totals. It selects an interval but does not change the spikes within it; an exact tie is resolved by the earliest start. Empty categories contribute zero to entropy, and the implementation uses ln(max(K, 2)) for the cluster normalization.

For population-burst visualization, spikes in the selected window were pooled in 1-ms bins and convolved with a unit-area discrete Gaussian kernel (σ = 5 ms, truncated at ±5σ). Values outside the window were zero-padded. The mean and population standard deviation were calculated over the full smoothed window. Each maximal contiguous run above mean + 3 SD was counted as one population burst, without merging or boundary expansion. These labels were not used for clustering, window selection, voice assignment, or MIDI construction. No local field potential criterion was applied, so they are not ripple detections.

### Pitch rank, velocity, and channel gain

Units within an instrument cluster received distinct MIDI keys in the oriented leaf order. Melodic keys were selected from the five-degree template associated with the actual instrument family. The code first used the stated instrument range and broadened the search to keys 0–127 if necessary; it rejected clusters exceeding that template’s available keys. A centered span of the required length plus up to four degrees was sampled at equally spaced rounded indices. The percussion kit instead used successive drum keys beginning at 35. No two units could share a (channel, key) pair.

Velocity and channel volume were reduced with increasing firing rate to limit the contribution of frequently firing units. These settings are not calibrated measures of perceived loudness. Let ℓ_i_ = log_10_(max(r_i_, 10⁻¹²)), and let u_i_ = (ℓ_i_ − min_j_ ℓ_j_) / (max_j_ ℓ_j_ − min_j_ ℓ_j_). The minimum and maximum are taken over all units, with a minimum denominator of 10⁻¹². Let R_k_ be the sum of unit firing rates in cluster k.

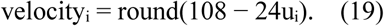

The attainable velocity range is 84–108.

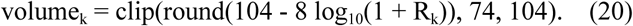

### Microtonal pitch systems

MIDI key numbers supplied an octave/register scaffold, whereas the custom SoundFont supplied a per-cell fine-tuning offset. D (pitch class 2) was the common tonic. The labels ’slendro-inspired,’ ’pelog-inspired,’ ’hirajoshi-inspired,’ and ’in-sen-inspired’ are mnemonic descriptions of the compositional palette, not claims of a canonical tuning or an ethnomusicologically authentic reconstruction. Tunings in these traditions vary by instrument, ensemble, region, and practice.

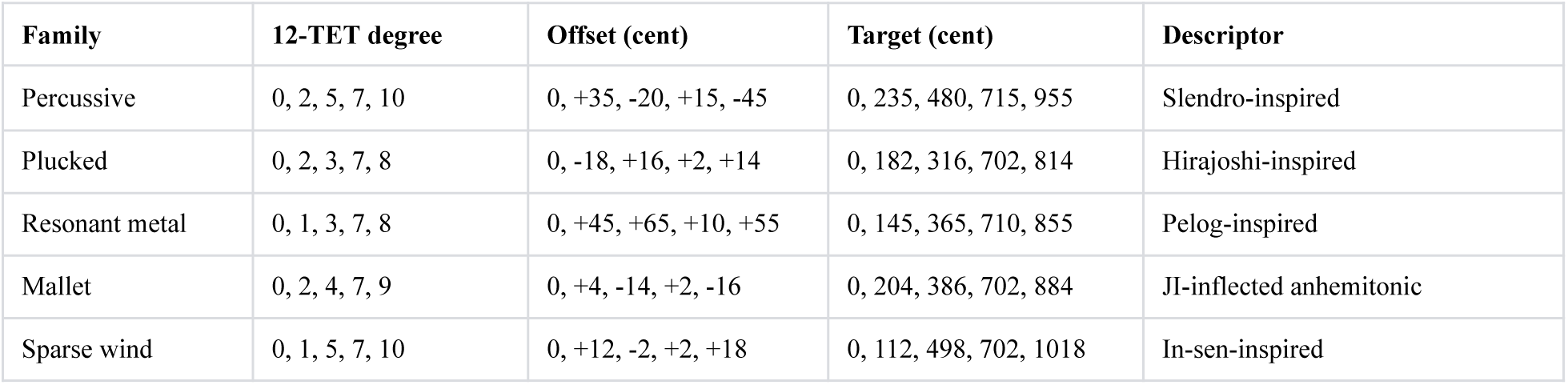

For a MIDI key q_i_ and fine-tuning offset c_i_ in cents, the nominal equal-tempered reference frequency is

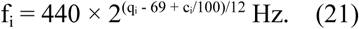

This formula assumes conventional key tracking and is not a measurement of the synthesized fundamental. The source bank may introduce additional tuning or inharmonic spectra. Percussion-kit samples receive offsets from the key-indexed cycle c = (−31, −12, +19, +43, −28, +8, +31, −17, +25, −42, +14, +37) cents. For these samples, a key selects a drum sound, and detuning shifts its spectrum without establishing the fundamental predicted by Eq. 21.

### Per-cell stereo spatialization

MIDI CC10 was held at its center value (64) on every channel. Because a channel controller applies to all units sharing that channel, separate pan settings were written into singleton-key SoundFont zones. In SoundFont units, −500 is full left, 0 is center, and +500 is full right. Values were clipped to [−480, +480].

Each instrument cluster k received a base pan β_k_ from the 16-channel template in the code. The channel-1 anchor and channel-10 percussion kit were centered. For rank a among n_k_ units, a branch coordinate was added with spread s_k_: 70 units by default, 90 for the centered melodic channel, and 170 for the percussion kit.

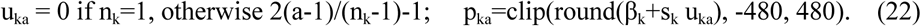

A normalized pan coordinate is *x_i_* = *p_i_*/500. An equal-power interpretation is *g*_L_ = cos[π(*x_i_*+1)/4] and *g*_R_ = sin[π(*x_i_*+1)/4]. The actual audio was generated by the SoundFont renderer; this expression states the intended monotonic spatial mapping rather than replacing the renderer’s panning implementation.

### Event-faithful MIDI clock and global time dilation

Let *q* be source-clock ticks per source second, *P* the Standard MIDI File division in ticks per quarter note, μ the tempo in microseconds per quarter note, *r* the desired playback rate relative to the source, and *t*_0_ the selected-window start. Each retained spike was converted once:

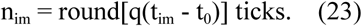

MIDI playback time at tick n is nμ/(10⁶P) seconds. For a tempo representable exactly as an integer number of microseconds per quarter note,

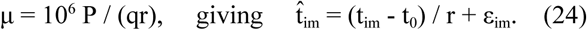

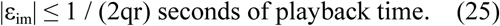

The defaults q = 30,000 ticks/s, P = 15,000 ticks/quarter, and r = 0.05 yield μ = 10,000,000 µs/quarter exactly. At these settings, Eq. 25 gives a maximum playback-time rounding error of 0.333 ms, equivalent to 16.7 µs in source time. For other rates, the code rounds μ to an integer; the effective rate is then r_eff = 10⁶P/(qμ), and Eqs. 24–25 apply with r_eff. Relative to the requested rate r, an additional drift term of at most W|1/r_eff − 1/r| must be included. The code rejects tempos outside the three-byte field (1–16,777,215); it does not automatically change P or implement a multi-tempo fallback.

A Note Off was placed at the earliest of the nominal duration cap, one tick before the next spike of the same unit, and the final tick, subject to a minimum duration of one tick. Events sharing a tick were ordered Note Off before Note On and then by key. Distinct input spikes can round to the same tick; the writer preserves their Note On count, but overlapping same-key messages may be interpreted differently by synthesizers. Thus the scheduling rule does not guarantee acoustic separation for arbitrary input timing. The final tick can also be extended by one tick when needed to retain the last onset.

### MIDI organization and continuous resonance

A type-1 Standard MIDI File was written with one conductor track and one track per instrument cluster. Each cell retained a unique (channel, key) pair within a one-port file. The melodic custom program was channel minus one; the General MIDI percussion preset used channel 10 and program 0. At tick zero, every active channel received bank selects, program change, channel volume, centered pan, sustain, reverb send, and chorus send.

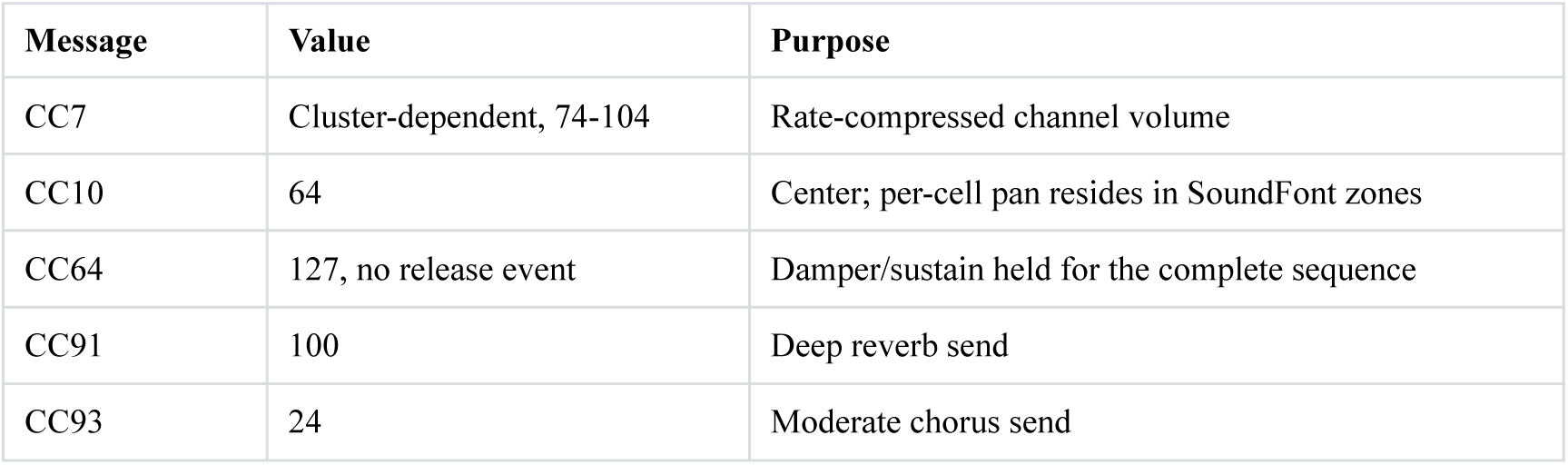

Sustain remained on through End of Track. This was a sound-design choice intended to retain overlapping resonances. For looped or sustaining samples, a Note Off need not produce an audible release while the pedal remains held. The four-second render extension is therefore an added tail interval, not a guarantee that every voice has decayed to silence.

### Custom SoundFont construction

The source SoundFont was loaded and cloned with spessasynth_core. The reported example used a bank identified as GeneralUser GS 2.0.3 BETA; the script copied its attribution and comment metadata into the derivative bank. The exact source bank, including its download source, digest, and applicable license text, is required to reproduce the timbres. Substituting another bank can change preset coverage, tuning, envelopes, and the resulting audio even when MIDI events remain unchanged.

For each instrument cluster, the source preset was cloned into a dedicated custom preset. Source zones covering a mapped MIDI key were copied into singleton key zones [q_i_,q_i_]. The cell’s cent offset was added to the copied zone’s existing fine tuning, and its pan generator was set to p_i_. The bank was trimmed to mapped keys and to each performed velocity plus representative layers 32, 64, 96, and 120; unused instruments and samples were removed. This operation changes storage and orchestration, not event content.

Global preset envelopes and effects were adjusted by instrument class. SoundFont timecents *tc* correspond to 2^(tc/1200)^ seconds, and absolute filter cents *fc* correspond approximately to 8.176 × 2^(fc/1200)^ Hz. Reverb and chorus generator values are in 0.1% units. The exact instrument-specific constants are included in the JavaScript listing.

| Shape | Attack $tc$ | Release $tc$ | Filter $fc$ | Reverb | Chorus |
| --- | --- | --- | --- | --- | --- |
| Deep drum | -12000 | -2100 | 10300 | 470 | 80 |
| Wind | -4700 | -500 | 11800 | 570 | 160 |
| Bright metal | -10000 | +650 | 13200 | 640 | 130 |
| Soft metal | -8200 | +250 | 12200 | 590 | 190 |
| Wood | -12000 | -1900 | 11200 | 410 | 60 |
| Soft wood | -10500 | -950 | 10700 | 440 | 90 |
| Drum kit | -12000 | -2300 | 11800 | 360 | 40 |
| Plucked | -11500 | -850 | 11600 | 500 | 100 |

### Offline synthesis, normalization, and MP3 encoding

The MIDI and derived bank were rendered offline with spessasynth_core 4.3.22 at 44,100 samples/s, using 128-sample synthesis blocks, enabled effects, and automatic voice allocation. The sequencer was advanced once per block, corresponding to about 2.90 ms of playback time. This block-based scheduling must be distinguished from the finer timing encoded in the MIDI file; sample-level onset alignment was not measured. Four seconds were appended after the nominal MIDI duration. The two floating-point channels were scaled by the same bounded gain,

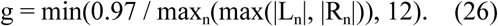

For a silent render, g was set to one. Applying the same gain to both channels preserves their relative amplitudes. Samples were bounded to [−1, 1] during 16-bit conversion and encoded as stereo, 44.1-kHz, constant-bit-rate 256-kbit/s MP3 using @breezystack/lamejs 1.2.7. The script does not save a lossless audio master. A PCM or lossless export should be retained for acoustic timing measurements, because MP3 encoding and playback can introduce additional delay or transient smearing.

### Verification and reproducibility criteria

Verification has three distinct levels: checks performed by the main scripts, independent tests of serialized MIDI, and tests still needed for the synthesized audio. A PASS status refers to the checks implemented by the relevant script; it is not evidence of complete scientific or perceptual validation.

- Event counts: the Python writer compares its Note On count with the selected spike count and compares per-unit counts after writing. These are internal writer checks, not an independent parse of the serialized file.
- Encoded times: smoke_test.py independently parses a synthetic-data MIDI file and compares the complete sorted list of (channel, key, tick) events with the selected input spikes. The main writer does not itself perform this readback check on every dataset.
- Voice addressing: the mapping must contain each input unit once, with no duplicate (channel, key) pair.
- Clock: the tempo must fit 24 bits. The effective rate is 10⁶P/(qμ); it equals the requested rate exactly for the reported default configuration.
- Controllers: the writer places CC64 = 127 and CC91 = 100 at tick zero, with the other controller values specified above. The supplied smoke test does not independently assert the controller state.
- Voice existence: the SoundFont script checks that every mapped (preset, key, velocity) yields at least one voice before serialization and after reload.
- Tuning and pan: the singleton-zone generator arrays are compared before serialization and after reload. This checks persistence of stored parameters, not the acoustic result of all combined generators and modulators.
- Stereo output: the script requires c_LR_ = Σn LnRn / √[(Σn Ln²)(Σn Rn²)] < 0.995 and normalized RMS(L − R) > 10⁻⁴. Here c_LR_ is an uncentered normalized inner product, not a mean-subtracted Pearson correlation. These thresholds reject nearly identical channels but do not measure spatial accuracy or source separability.
- Provenance: the Python manifest records Config values, recording and excerpt bounds, counts, effective cluster capacity, tempo, and SHA-256 digests of the source CSV and MIDI. The audio manifest records source-bank, derived-bank, and MP3 digests plus selected render parameters and stereo metrics. Software versions are specified in the dependency files rather than comprehensively captured at runtime.
- Reproduction: retain both manifests, the mapping CSV, the exact source and derived banks, code revision, dependency lockfiles, and runtime versions. Several mapping constants and tie rules reside in the source code rather than the manifest. The renderer inserts a new creation date into the derived bank, so repeated builds need not have identical bank digests even when the mapping is unchanged.

The supplied Python smoke test exercises the pipeline on a synthetic raster with 40 units and independently checks the serialized Note On events. The mathematical test checks selected numerical identities and parameter ranges; it does not validate every numbered equation or the full clustering procedure. The reported SoundFont checks concern voice existence, stored tuning and pan, and stereo output. Neither test establishes that the synthesizer avoids voice stealing or that every retained spike produces a separately detectable acoustic onset.

### Parameter scaling and boundary conditions

Storage is dominated by the B × N filtered arrays and N × N dissimilarity matrices, with additional costs for hierarchical and dynamic-programming calculations. Chunked covariance estimation, sparse graphs, and multiple MIDI ports are possible extensions, but are not implemented in the supplied reference code. The current implementation assumes 2 ≤ G ≤ K ≤ 16, positive analysis and excerpt parameters, and sufficient available keys in every assigned instrument. Percussion-kit addressing is limited to the 47 keys from 35 through 81; larger clusters require an explicit capacity check or a revised mapping. Validation of these limits should precede use on substantially larger inputs.

The 12 resampling replicates provide an economical co-association term for orchestration. They are not a confidence-interval procedure, and no improvement in stability was measured against an unresampled control. Family names, instrument pools, and tuning templates are compositional choices rather than claims about neural cell classes or authentic reconstructions of particular musical traditions.

### Software, standards, and source resources

- The MIDI Association. Standard MIDI Files and MIDI 1.0 specifications. https://midi.org/standard-midi-files
- Spessasus. spessasynth_core 4.3.22 documentation. https://spessasus.github.io/spessasynth_core/
- Collins SC. GeneralUser GS 2.0.3, License v2.0. https://schristiancollins.com/generaluser.php
- Creative Technology Ltd. SoundFont 2.04 Technical Specification. https://www.synthfont.com/sfspec24.pdf
- The code was developed with assistance from gpt-5.6-sol; further disclosure is provided in the Acknowledgments.

### Animal experiments

All experiments were performed with the approval of the Animal Experiment Ethics Committee of the University of Tokyo (approval number P29-7) and in accordance with the University of Tokyo guidelines for the care and use of laboratory animals. A male Long–Evans rat (6 months old, approximately 400 g at the time of surgery) was purchased from Japan SLC (Shizuoka, Japan) and used in this study. The animal was housed individually and maintained on a 12-h light/12-h dark cycle with lights off at 7:00 A.M. For electrode implantation, the rat was anesthetized with isoflurane gas (0.5–2.5%) and placed on a flat heating pad in a stereotaxic frame. Veterinary ointment was applied to the eyes to prevent drying during anesthesia, and buprenorphine (0.05 mg/kg, s.c.) was administered as an analgesic. A 2-cm midline incision was made in the scalp between the eyes and the cerebellum to expose the skull. Two craniotomies, each 1.5 mm in diameter, were made bilaterally above the dorsal hippocampus (3.5 mm posterior and 3.3 mm lateral to bregma) using a high-speed dental drill, and the dura mater was carefully removed. Two silicon probes (H32, A-style; NeuroNexus, Ann Arbor, MI, USA) were stereotaxically positioned above the craniotomies, lowered to the cortical surface, and inserted 1.0 mm into the brain at the end of surgery. All recording devices were secured to the skull with stainless-steel screws and dental cement. A screw placed above the cerebellum served as the ground and reference electrode. After surgery, the rat was housed individually in a transparent Plexiglas cage with ad libitum access to food and water for 7 d. Following recovery, the animal was food-restricted until its body weight reached 85% of the preoperative value. During this period, the probes were advanced in steps of 100 µm per day until the recording sites reached the CA1 pyramidal cell layer, as identified by the presence of sharp-wave ripples and unit activity.

Neural activity was recorded continuously from dorsal CA1 for 61 min while the rat freely explored its home cage. Signals were amplified and digitized at 30 kHz using an Open Ephys recording system and stored for offline analysis. Before spike sorting, the broadband signals were band-pass filtered (300–6000 Hz) and common-average referenced across channels to attenuate shared noise. Spikes were sorted offline using Kilosort4 (Pachitariu et al., 2016, 2024), with default parameters except where noted. The resulting clusters were inspected and manually curated in Phy (Rossant et al., 2016). Curation considered waveform shape and consistency across channels, a clear refractory period in the autocorrelogram (violations within ±2 ms below 0.5% of all interspike intervals), stable spike amplitudes throughout the recording, and separation from neighboring clusters in feature space. Clusters with multi-unit characteristics, amplitude drift, or refractory-period contamination were merged, split, or discarded as appropriate. Each retained cluster was judged to represent a single, well-isolated unit.

Each well-isolated unit was classified as a putative pyramidal cell or interneuron from its extracellular spike waveform. The mean waveform was calculated on the channel with the largest spike amplitude, and waveform width was measured from the negative trough to the subsequent positive peak (trough-to-peak latency). Extracellular waveform width can help distinguish principal cells from interneurons (Csicsvari et al., 1999; Barthó et al., 2004), although it does not establish cell identity. The observed distribution of trough-to-peak latencies was bimodal. Units with a latency shorter than 0.5 ms were classified as putative interneurons and excluded. The remaining 168 putative pyramidal units formed the dataset used for sonification.

## Results

### Dataset

The event table contained 168 putative CA1 pyramidal-unit identifiers and 1,501,913 spikes over a 3,661-s recording (approximately 61 min). Acquisition was reported at 30 kHz; this information and the unit classifications derive from the experimental record, not from independent fields in the CSV. Mean unit firing rates ranged from 0.013 to 9.7 Hz, with a median of 1.7 Hz. The reference code classified 732,652 of 1,501,745 within-unit ISIs (48.79%) as shorter than 50 ms after conversion to floating-point seconds. The small boundary-dependent difference from a millisecond-valued calculation is described in Methods. The full recording informed the voice mapping; the event output comprised the selected excerpt.

### Orchestration derived from population structure

The derived partition grouped units with lower average dissimilarity within instrument clusters (0.80) than between clusters (0.95). Mean bootstrap co-association was 0.60 within instrument clusters, 0.48 between clusters in the same supercluster, and 0.08 between superclusters. At τ = 0.1 s, mean correlation was 0.18 within clusters and 0.07 between them; at τ = 3 s, the corresponding means were 0.35 and 0.07.

These values describe the partition obtained from the same data and dissimilarity used to construct it. They are not independent evidence of discrete biological assemblies or of out-of-sample cluster stability.

The partition contained five superclusters of 16, 1, 101, 22, and 28 units, divided into 16 instrument clusters of 1–15 units at meff = 15 (Figure 2). The percussive supercluster contained 16 units and had a median rate of 7.2 Hz; 14 units received the GM percussion kit and two received taiko drum. The singleton supercluster, firing at 0.013 Hz, received shakuhachi. The 101-unit supercluster had a median rate of 0.59 Hz and received the resonant-metal family label. Its instruments included five resonant-metal presets carrying 61 units, agogo carrying 13 units from the family pool, and fallback woodblock and melodic tom presets carrying 13 and 14 units. The remaining superclusters received mallet (22 units; median rate 3.9 Hz) and plucked (28 units; median rate 3.1 Hz) assignments. The five supercluster labels therefore differ from the families of the final instruments: by actual instrument family, the mapping contained 56 percussive, 61 resonant-metal, 22 mallet, 28 plucked, and one wind unit. The mean-based assignment rules are specified in Methods; the medians reported here summarize the groups.

**Figure 2.**
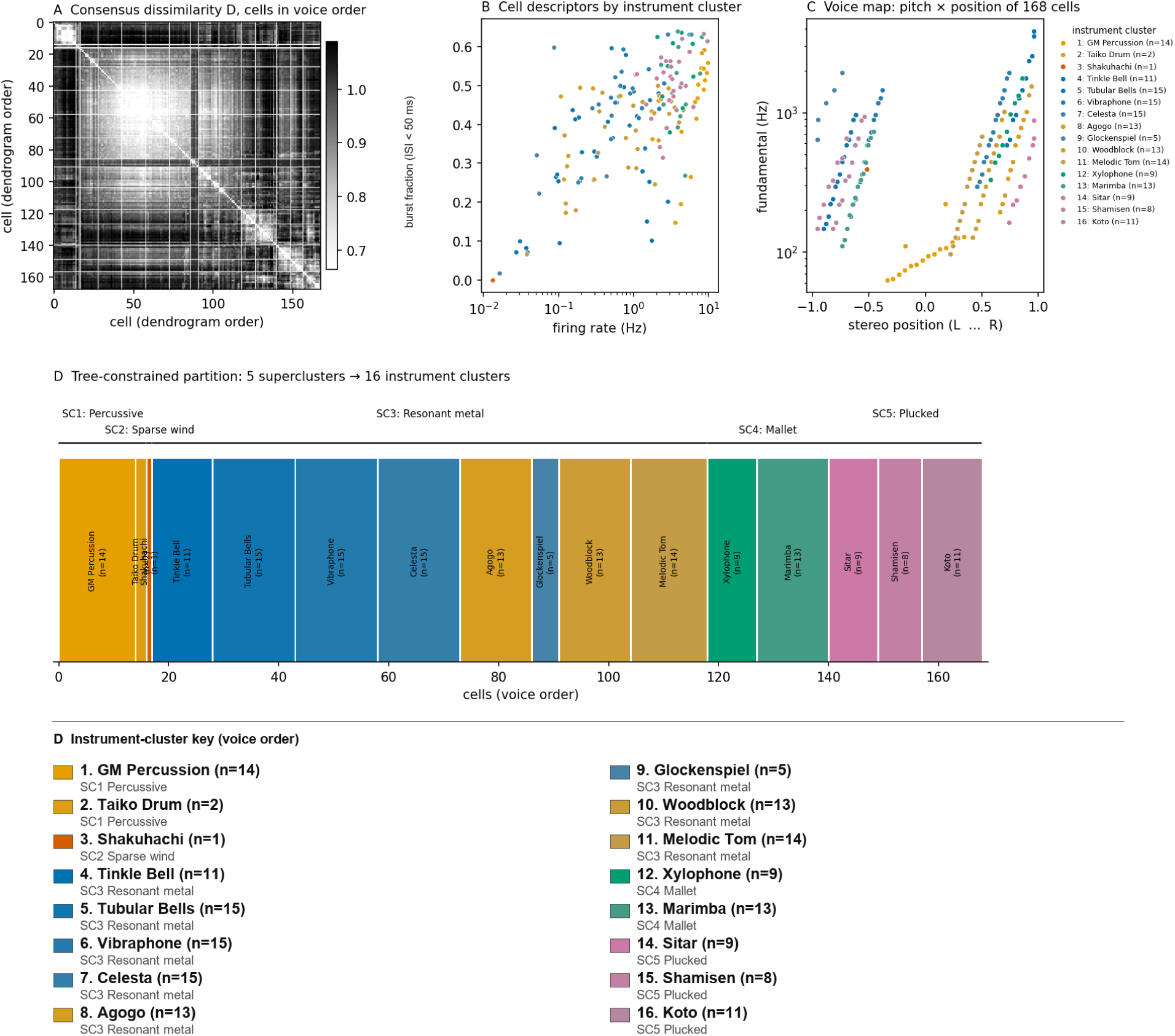
Orchestration of the recorded population. (A) Combined dissimilarity D among 168 unit identifiers, shown in voice order. White lines mark instrument-cluster boundaries. (B) Firing rate and short-ISI fraction (ISI < 50 ms), colored by actual instrument family and cluster. (C) Pan coordinate (SoundFont pan divided by 500) and nominal reference frequency calculated from key and offset using Eq. 21. The plotted frequencies are not measured acoustic fundamentals, particularly for percussion-kit keys. (D) Five superclusters and their 16 instrument clusters. The key beneath the partition lists each instrument, unit count, parent supercluster, and assigned supercluster family. The term cell in the panel labels refers to an input unit identifier.

Every unit had a distinct (channel, key) pair. Frequencies calculated from keys and offsets using Eq. 21 ranged from approximately 63 to 3,850 Hz, offsets ranged from −45 to +65 cents, and pan settings ranged from −480 to +480 SoundFont units (Figure 2C). These are nominal reference frequencies, not measured fundamentals, particularly for the percussion kit. Velocities ranged from 84 to 108 and channel volumes from 88 to 104. The 24-unit velocity range cannot be interpreted as a loudness ratio, because the amplitude response depends on the instrument and velocity layer.

### The sonified excerpt

The window score selected 3,590–3,620 s, containing 12,219 spikes from 166 of the 168 units (1–328 spikes among active units; median 63). At r = 0.05, the nominal duration is 600 s, followed by the four-second render extension. The population-count threshold identified 44 contiguous suprathreshold epochs; no local field potential was used to establish ripple identity. The densest 128-ms interval contained 316 spikes from 99 units and occupies 2.56 s at this dilation (Figure 3B, C). Dividing that duration by the event count gives about 8.1 ms per event, but this average does not specify the minimum separation or establish that individual events are perceptually resolvable. The excerpt also contained 2,269 within-unit ISIs shorter than 6 ms, corresponding to less than 120 ms in playback. This additional interval count describes rapid firing; it is neither a validated perceptual threshold nor the 50-ms descriptor used for clustering.

**Figure 3.**
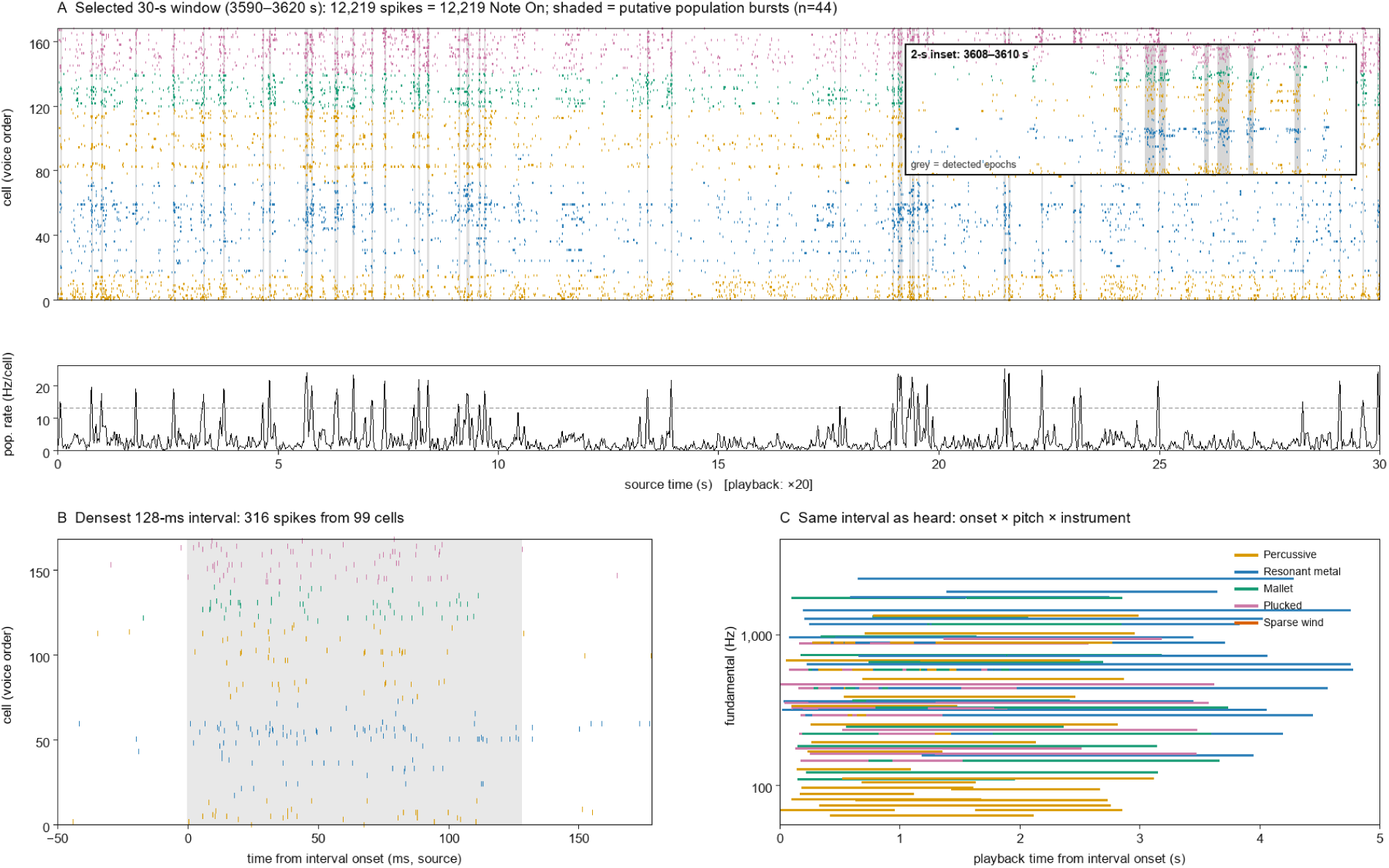
Selected excerpt and event representation. (A) Raster of 3,590–3,620 s, arranged in voice order and colored by instrument cluster. Shading indicates 44 contiguous population-count epochs above mean + 3 SD after 1-ms binning and Gaussian smoothing (σ = 5 ms). The population trace below is expressed as a rate per recorded unit; the dashed line marks the threshold. The inset enlarges 3,608–3,610 s. Relative source time zero is 3,590 s, and one source second corresponds to 20 playback seconds. (B) The densest 128-ms interval, containing 316 spikes from 99 units, with surrounding activity shown for context. (C) A score-like representation using playback onset and nominal reference frequency. This panel depicts the assigned events, not measured acoustic onsets or fundamentals. Horizontal spans illustrate the nominal duration settings; actual Note Off times may be shortened before a subsequent spike, and sustain and effects can extend audible decay.

### Verification of event fidelity

Independent reproduction of the supplied implementation yielded 12,219 Note On messages for 12,219 selected spikes, with matching per-unit counts (Figure 4A). Independent parsing gave a maximum source-time deviation of 0.33 µs, below the 16.7-µs bound in Eq. 25 after conversion to source units (Figure 4B). This small nonzero deviation is consistent with timestamps lying close to, rather than exactly on, the 30-kHz clock grid. The stored P = 15,000 and μ = 10,000,000 µs/quarter give r_eff = 0.05 exactly. Encoded interspike intervals follow twentyfold dilation within the rounding error of their two endpoint times (Figure 4C). Input and MIDI digests are listed in Table 1. These results concern encoded MIDI events, not measured acoustic onset times.

**Figure 4.**
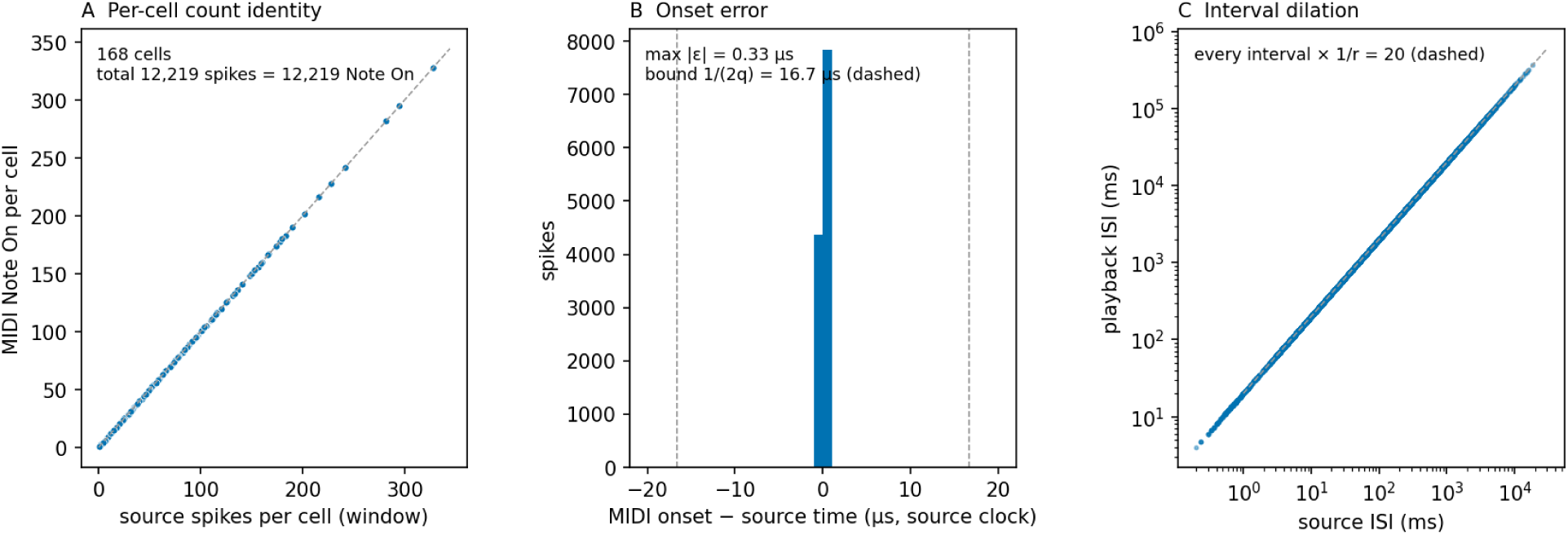
Verification of encoded events. (A) Source-window spike count versus MIDI Note On count for each unit. (B) MIDI onset error expressed in source-time units; dashed lines show ±1/(2q). (C) Encoded playback ISIs versus source ISIs; the dashed line shows the nominal dilation factor 1/r = 20. The finite onset errors in B also bound deviations of intervals in C. These panels assess the MIDI event representation, not acoustic onset detection.

**Table 1.**
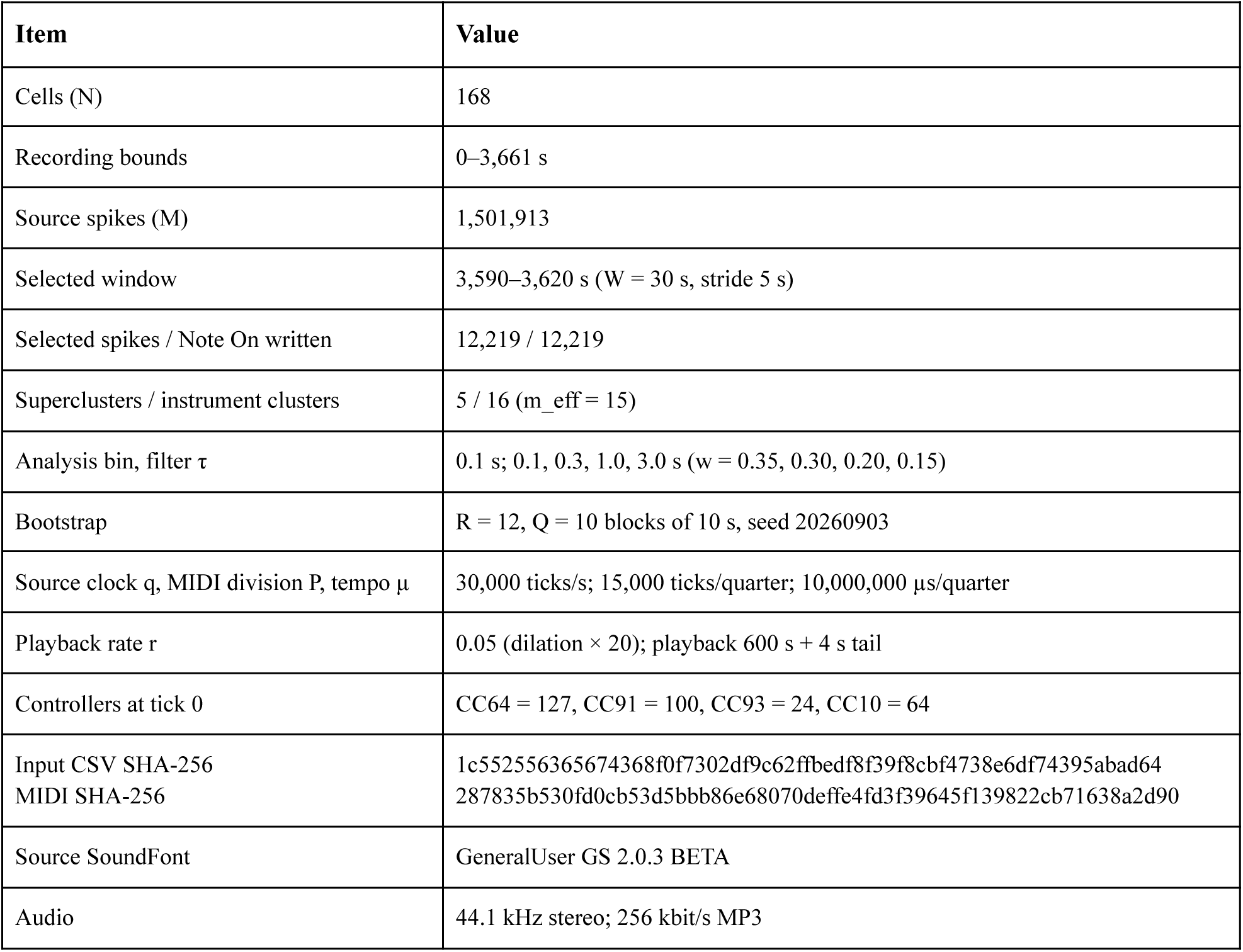
Manifest of the rendered example.

| Item | Value |
| --- | --- |
| Cells (N) | 168 |
| Recording bounds | 0–3,661 s |
| Source spikes (M) | 1,501,913 |
| Selected window | 3,590–3,620 s (W = 30 s, stride 5 s) |
| Selected spikes / Note On written | 12,219 / 12,219 |
| Superclusters / instrument clusters | 5 / 16 ($m_{\text{eff}} = 15$) |
| Analysis bin, filter $\tau$ | 0.1 s; 0.1, 0.3, 1.0, 3.0 s ($w = 0.35, 0.30, 0.20, 0.15$) |
| Bootstrap | R = 12, Q = 10 blocks of 10 s, seed 20260903 |
| Source clock q, MIDI division P, tempo $\mu$ | 30,000 ticks/s; 15,000 ticks/quarter; 10,000,000 μs/quarter |
| Playback rate r | 0.05 (dilation × 20); playback 600 s + 4 s tail |
| Controllers at tick 0 | CC64 = 127, CC91 = 100, CC93 = 24, CC10 = 64 |
| Input CSV SHA-256<br>MIDI SHA-256 | 1c552556365674368f0f7302df9c62ffbedf8f39f8cbf4738e6df74395abad64<br>287835b530fd0cb53d5bbb86e68070deffe4fd3f39645f139822cb71638a2d90 |
| Source SoundFont | GeneralUser GS 2.0.3 BETA |
| Audio | 44.1 kHz stereo; 256 kbit/s MP3 |

### Listening

An online listening example is available at https://www.youtube.com/watch?v=zOYGXcp727A. In informal listening, I heard population bursts as brief increases in ensemble activity and could follow some changes in the order and participation of voices. Rapid within-unit firing produced repeated notes on a fixed instrument, particularly in the plucked and mallet sections. Lower-rate units assigned to resonant instruments produced a more sustained background, and the singleton shakuhachi unit had two Note On events in the excerpt. Note On times derived from the spikes, whereas pitch contours, timbral grouping, apparent prominence, and acoustic decay also depended on the chosen mapping and renderer.

## Discussion

HERO combines event-based neural sonification with a specified procedure for organizing a large ensemble of recorded units. Distinct tones or instruments for neuronal sources and onset-preserving mappings have clear precedents (Aertsen and Erb, 1987; Hermann et al., 2002; Baier, Hermann and Stephani, 2007; Destexhe and Foubert, 2018, 2022). Its contribution is the integration of a fixed unit-to-voice mapping, hierarchical orchestration, fine-grained MIDI timing, and explicit verification procedures in one reference implementation.

The design is a form of parameter-mapping sonification (Kramer et al., 1999; Hermann, Hunt and Neuhoff, 2011; Dubus and Bresin, 2013). It addresses practical questions raised by multineuronal recordings: how to assign many voices consistently, how to avoid changing onset times during analysis, and how to retain enough information to audit the result.

The strongest fidelity statement concerns the encoded event list. A source timestamp is converted to a MIDI tick by one documented rounding operation, and the resulting events can be matched back to the input by unit, key, and time. The example and synthetic smoke test support this correspondence for the tested inputs. An acoustic waveform is a further transformation: sample attacks, block scheduling, sustained envelopes, effects, and encoding can alter the timing and salience of audible features. A rhythm perceived in the audio should therefore be checked against the source raster rather than treated as direct evidence of a neural sequence.

Separating analysis from event construction prevents the 100-ms analysis bins from becoming an unintended onset grid. Filtered activity controls the mapping, while original timestamps control the event list. The separation also makes controlled comparisons possible: alternative orchestrations can present the same spikes at the same encoded times.

The hierarchy provides a rule for organizing units with correlated activity into instrument groups. Because the analysis uses time constants of 0.1–3 s, it primarily describes co-modulation at those scales and does not resolve ripple-cycle timing or demonstrate synaptic connectivity. The family rules then translate selected descriptors into musical attributes. These choices are explicit but remain choices made by the designer; neither the clustering nor the palette establishes how listeners will group the sounds. Instrument-pool fallbacks also mean that some units receive an instrument family different from their parent supercluster label.

Singleton-key SoundFont zones allow units on a shared MIDI channel to retain separate tuning and pan settings. This extends per-unit control beyond what channel-level controllers alone provide, within the available key and channel limits. Fixed velocities and channel volumes reduce gain as firing rate rises, but the mapping has not been calibrated for loudness or tested for masking. Distinct voice addresses should not be equated with perceptually distinct sources.

Reproducibility requires more than a random seed. The exact source bank, realized mapping, code and numerical environment, and render settings all matter. The current manifests preserve useful provenance, but do not contain every constant, runtime version, or acoustic verification result. Preserving a tested release and its original artifacts is therefore necessary for exact audit, especially when a later numerical environment could change an ambiguous spectral ordering.

### Limitations

No listener study was performed, and the demonstration comprises one recording from one animal. It establishes an implementation example, not a general estimate of detectability or biological variation. Twentyfold dilation makes short intervals longer but also stretches slower fluctuations beyond their original timescale. Parvizi et al. (2018) tested whether listeners could classify sonified EEG samples, providing a relevant model for perceptual validation. For HERO, blinded comparisons could measure population-burst detection, sequence-order judgments, and source participation against source-data labels while holding event times fixed across orchestrations.

Sustain, long sample envelopes, reverb, and chorus spread energy over time and can promote perceptual fusion. Automatic voice allocation does not demonstrate the absence of voice stealing or interactions between percussion samples with exclusive-class behavior. The stereo check tests channel difference, not preservation of every unit’s spatial position. Measuring isolated and overlapping acoustic responses would be needed to characterize these effects and to distinguish MIDI fidelity from fidelity of the rendered waveform.

The analysis and orchestration parameters were chosen for this demonstration. They include bin width, filter scales and weights, resampling settings, λ, cluster counts and capacities, descriptor rules, and the instrument and tuning palettes. No ablation or sensitivity analysis establishes an optimum or demonstrates that hierarchy-based assignment improves listening performance. The co-association term summarizes repeated partitions of sampled trace segments; it should not be interpreted as confidence in a biological grouping. Likewise, the contribution of microtonal offsets to source segregation remains untested.

Assigned pitch encodes rank in the chosen unit order, not a physical property such as anatomical position or intrinsic firing frequency. Similarity-based ordering can generate melodic contours that listeners interpret as intentional sequences. Such impressions require comparison with the original timestamps and should not be assigned biological meaning from their musical form alone.

The result depends on spike sorting and curation. A neuron split across identifiers can appear as multiple voices, and contamination can combine spikes from different neurons into one voice. Split units may partition a spike train or duplicate some events; they do not necessarily sound in unison. Some units in this type of dataset were reported to be split across identifiers. Exact duplicate-row rejection does not resolve duplicate units.

One MIDI port limits the reference implementation to 16 instrument channels and the available keys within each instrument. Larger populations require changes to voice addressing. Auditory capacity imposes a separate limit: simultaneous sources may fuse or mask one another despite distinct keys, timbres, and pan settings (Bregman, 1990). No claim is made that listeners can track all 168 units independently.

The excerpt score favors participation, entropy, and temporal contrast. The selected 30 s should therefore not be treated as an unbiased sample of the full recording. Rendering the full event sequence avoids this selection step but produces a duration of approximately T/r, which is about 20.3 h for the present recording at r = 0.05.

The input representation contains spike times and unit identifiers only. It provides no waveform, amplitude, subthreshold, or field-potential information. In contrast, motif-based EEG sonification can map waveform morphology to sound envelopes (Destexhe and Foubert, 2022). Additional input streams would be needed to convey those measurements. In particular, spike-time sonification alone cannot establish the presence of a ripple oscillation or the replay of a previously measured behavioral sequence.

### Scope

HERO is a sonification method and reference implementation for exploring the temporal structure of spike trains. It preserves the selected event list within a documented MIDI clock conversion and makes the orchestration sufficiently explicit to inspect and test. The hippocampal example motivates perceptual experiments on population bursts, unit participation, and rapid firing. Establishing an advantage over simpler mappings will require those experiments and sensitivity analyses with the event sequence held fixed.

## Acknowledgments

I thank Nobuyoshi Matsumoto, Ryosuke Koike, and Ayako Ishigaki for assistance with hippocampal recordings. I used claude-fable-5-1 and gpt-6-astra for assistance with manuscript drafting and revision, and gpt-5.6-sol for code generation. Figure 1 was generated with gpt-image-2 and edited manually. I reviewed the resulting material and take responsibility for the manuscript and code.

## Author Contributions

Yuji Ikegaya conceived and designed the study, acquired and analyzed the data with the recording assistance acknowledged above, implemented the software, and wrote and revised the manuscript.

## Competing Interests

The author declares no competing interests.

## Funding

This work was supported by JST ERATO (JPMJER1801), JST Moonshot R&D (JPMJMS2012), the Institute for AI and Beyond of the University of Tokyo, JSPS KAKENHI (22K21353), and AMED Brain/MINDS 2.0 (JP24wm0625207, JP24wm0625401, JP24wm0625502).

## Code and data availability

Source event data and reference code are available in the HERO repository ( https://github.com/etangval/HERO). The implementation used for verification corresponds to repository commit 028c151. The appendix reproduces the implementation with typographical corrections and a more specific audit success message. The online audio is provided for listening; reproducibility requires the original rendered artifact and its digest.

## Appendix. Reference implementation

The following listings contain the reference files. Save each under its stated filename and run commands from that directory. The Python program uses the input unit count and recording duration, subject to the clustering and one-port addressing limits described in Methods. requirements.txt and package.json pin direct dependencies; reproducing the environment also requires runtime versions and dependency lockfiles. The examples require a Python version compatible with NumPy 2.3.2 and pandas 2.3.2, and a Node.js version compatible with spessasynth_core 4.3.22.

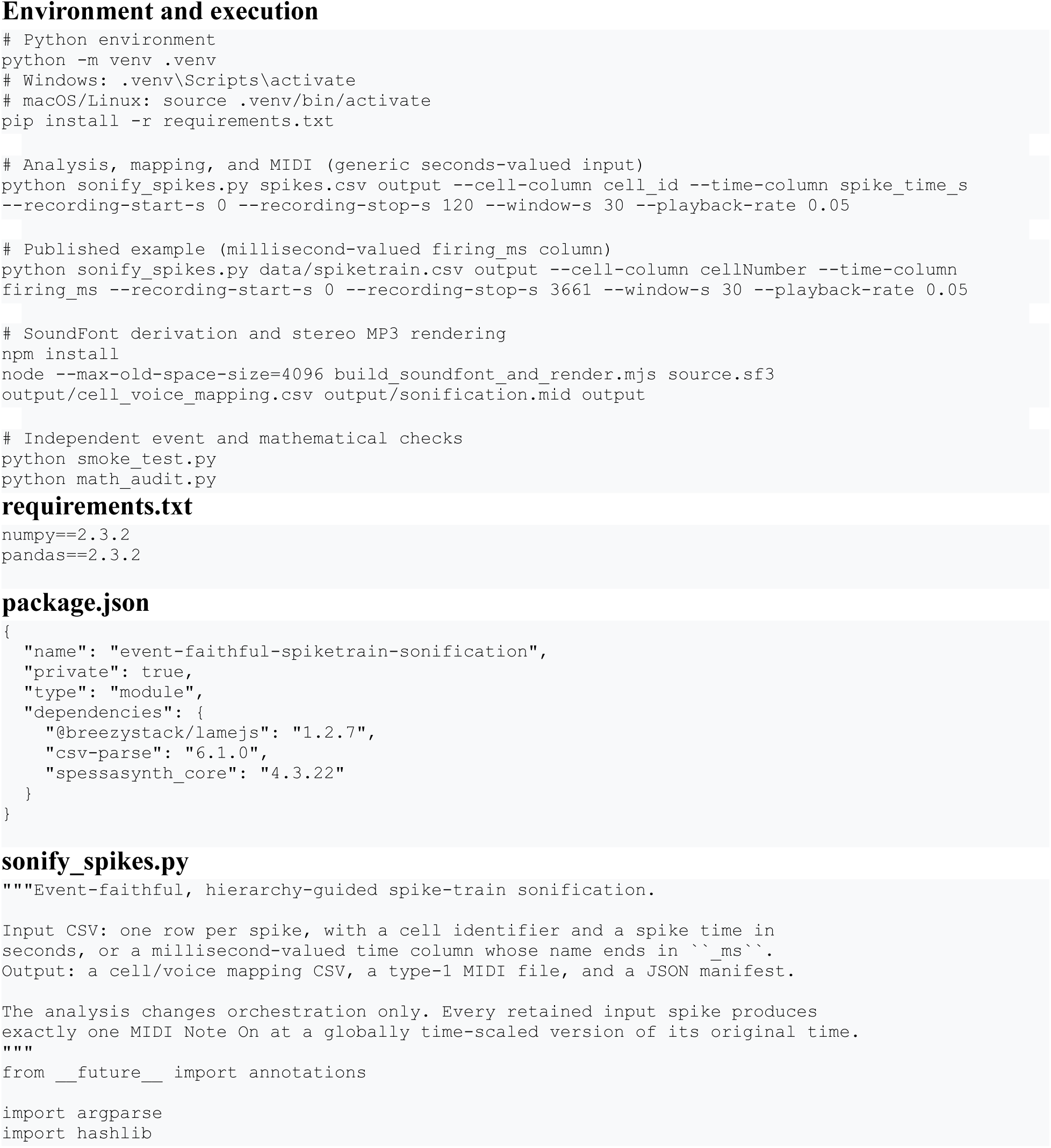
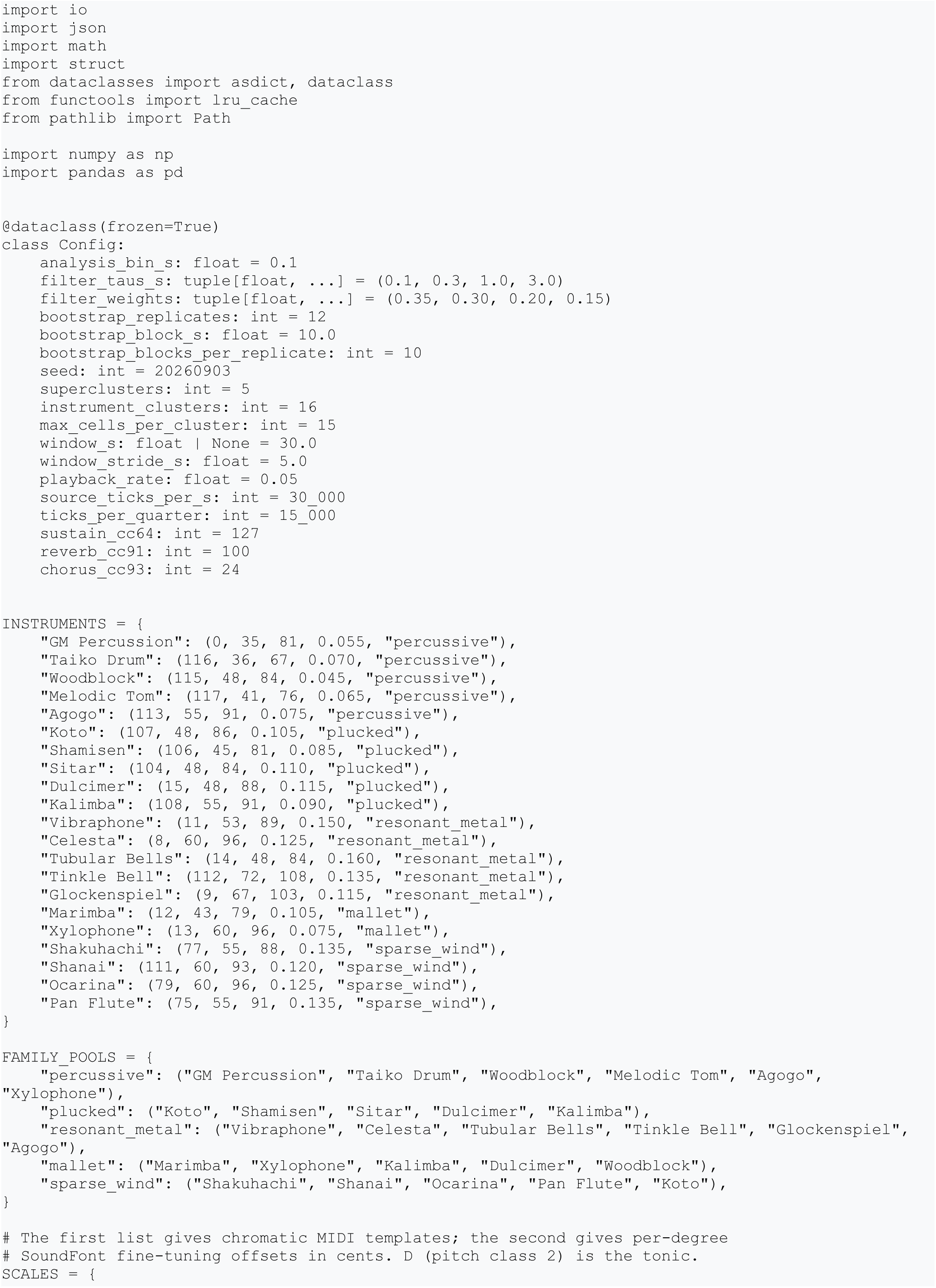
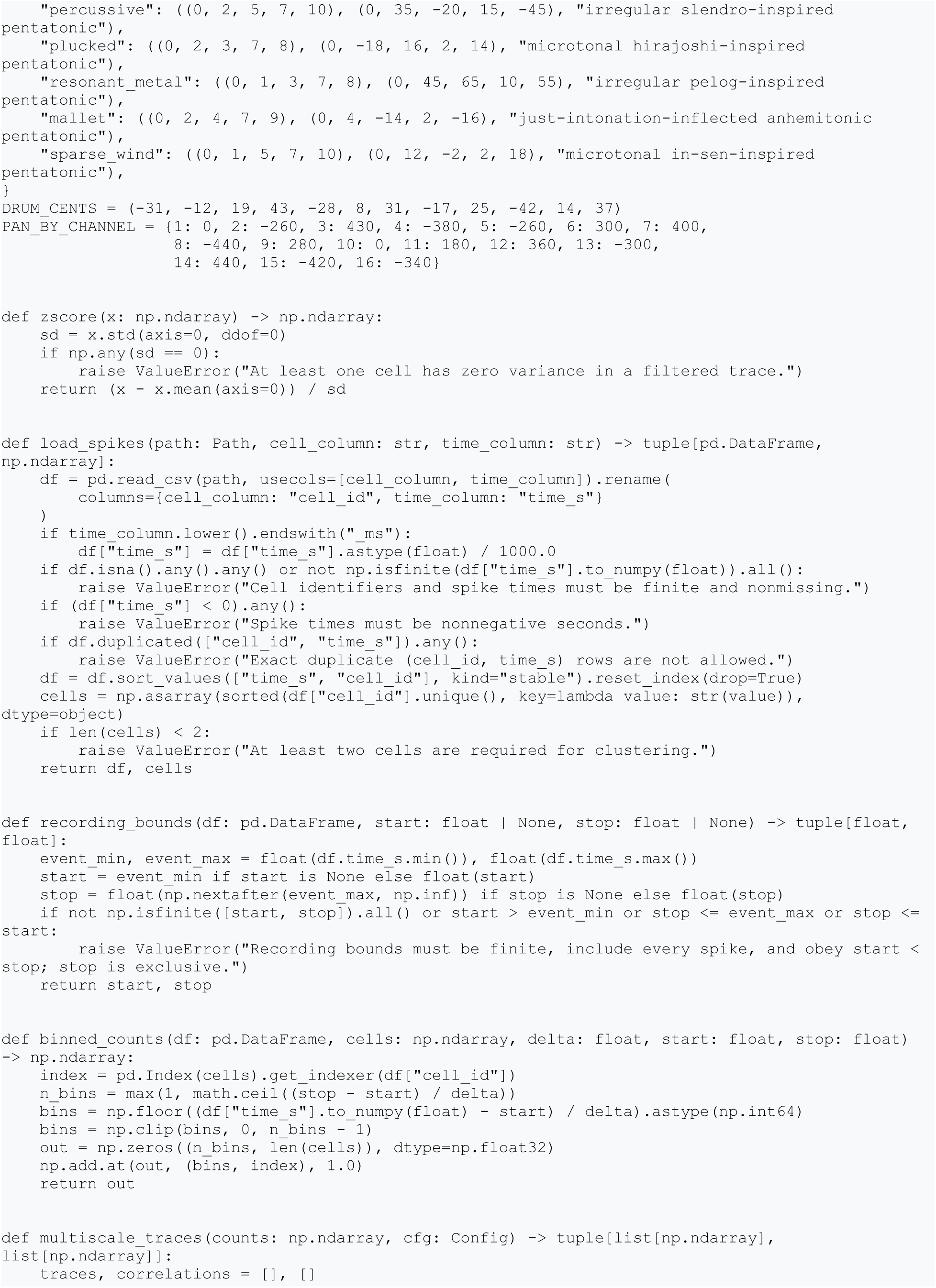
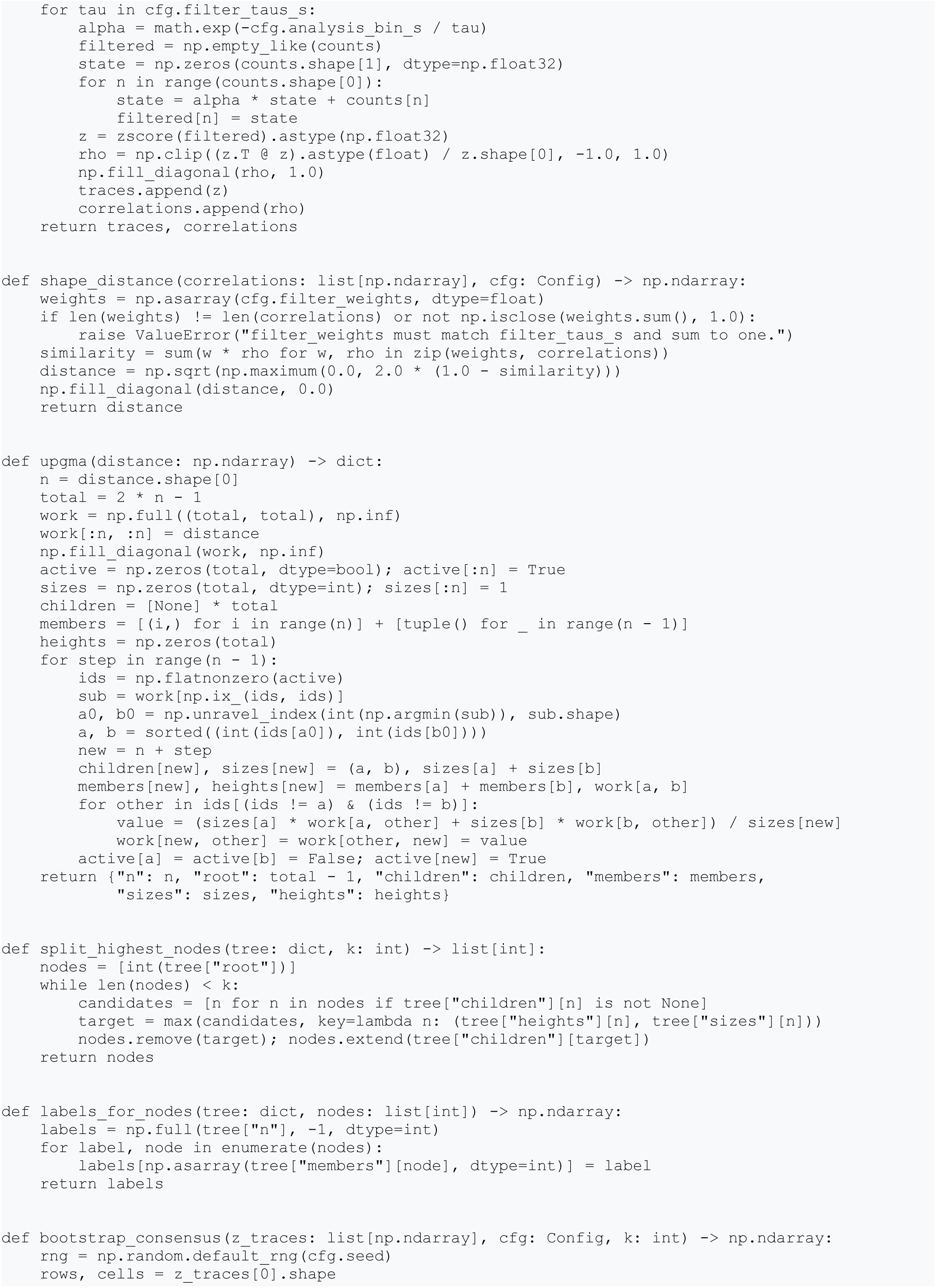
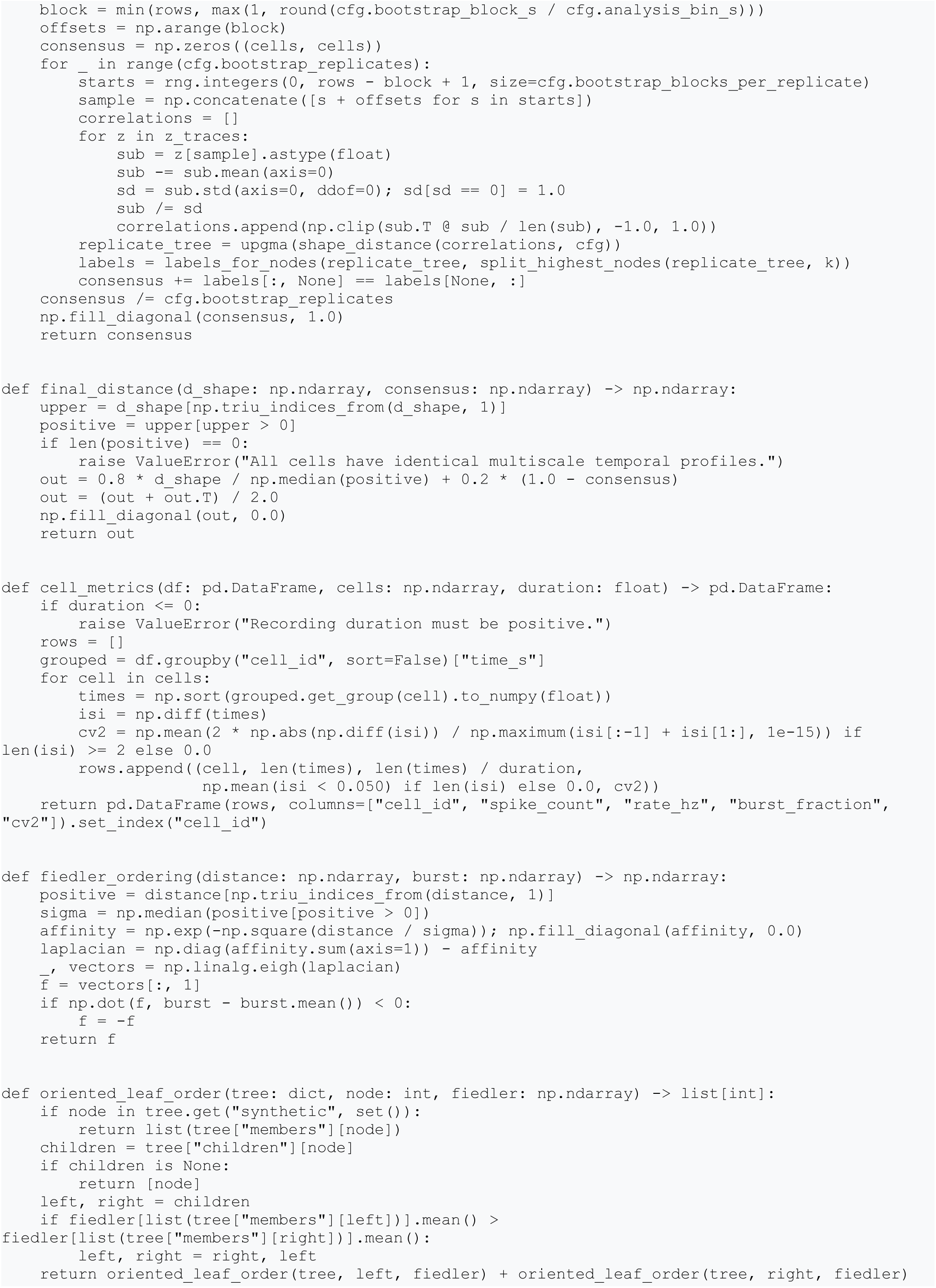
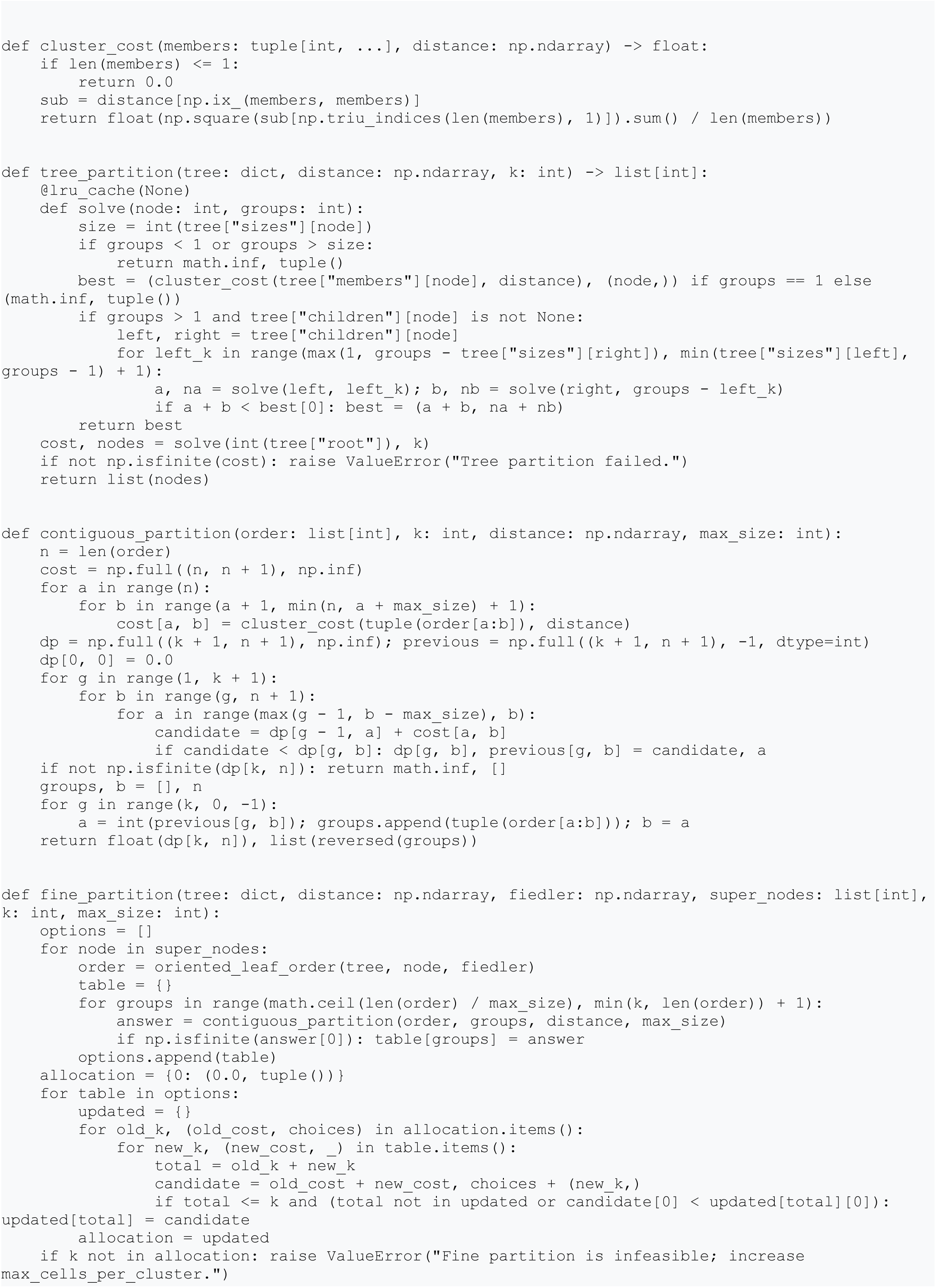
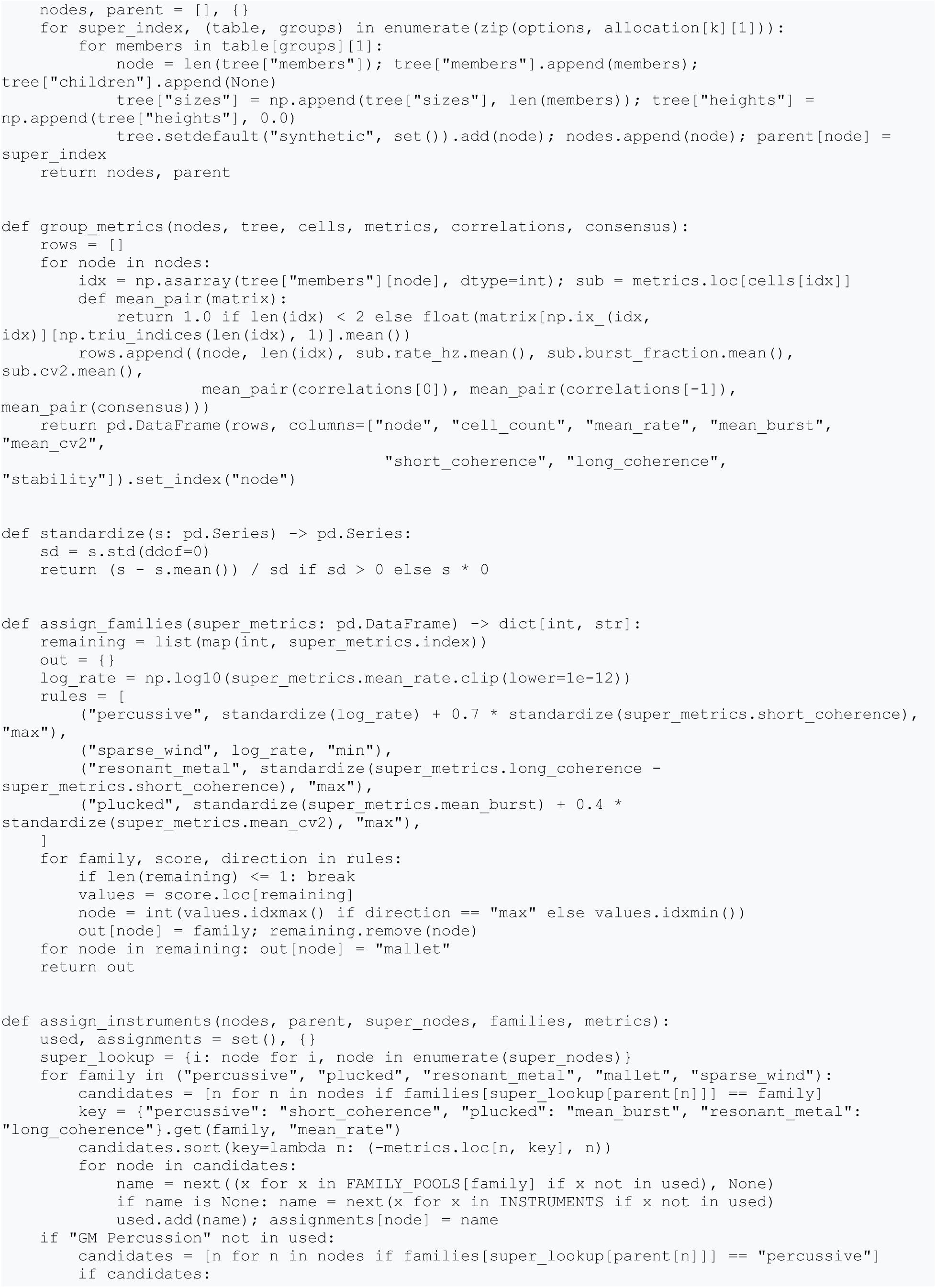
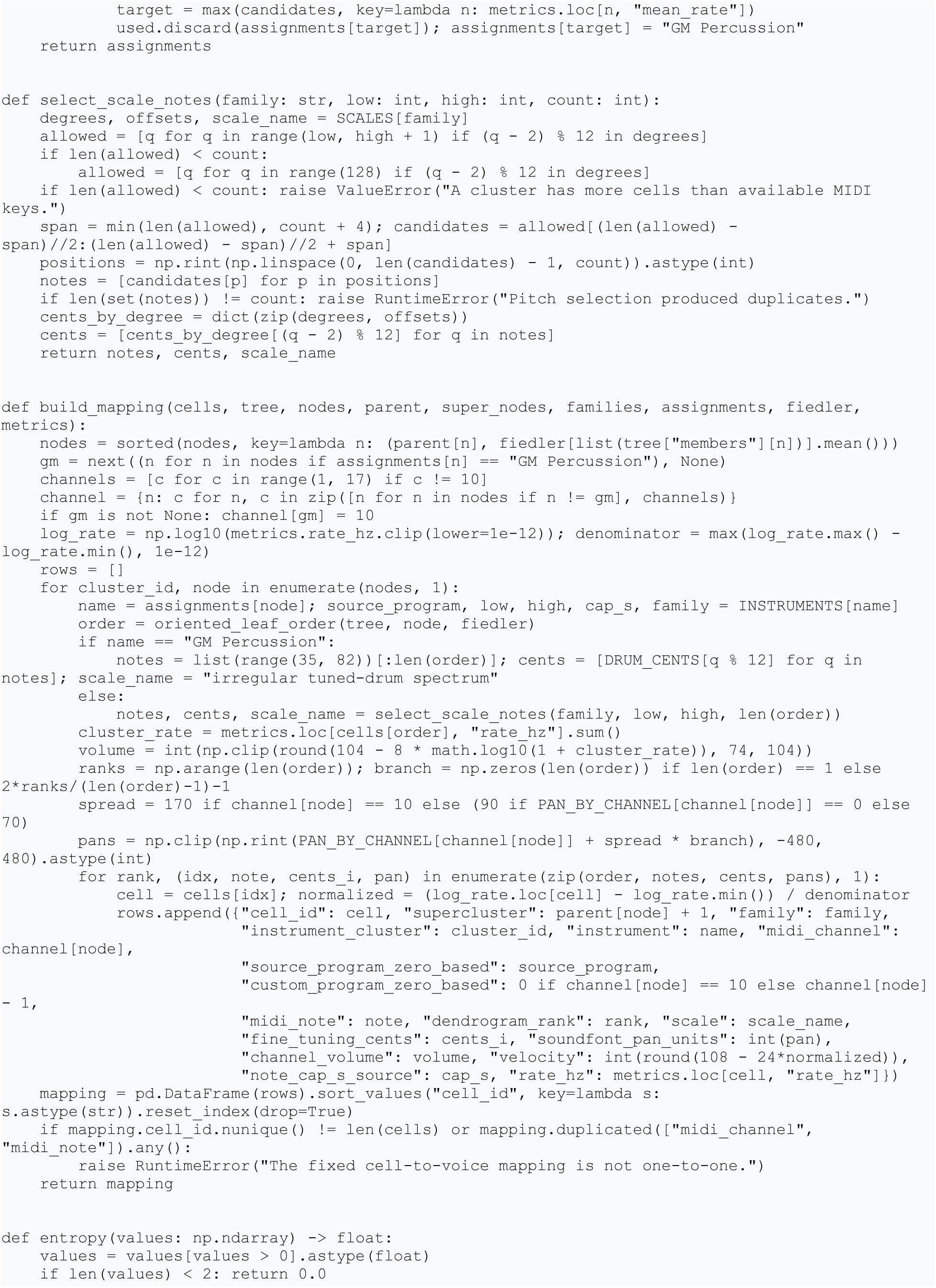
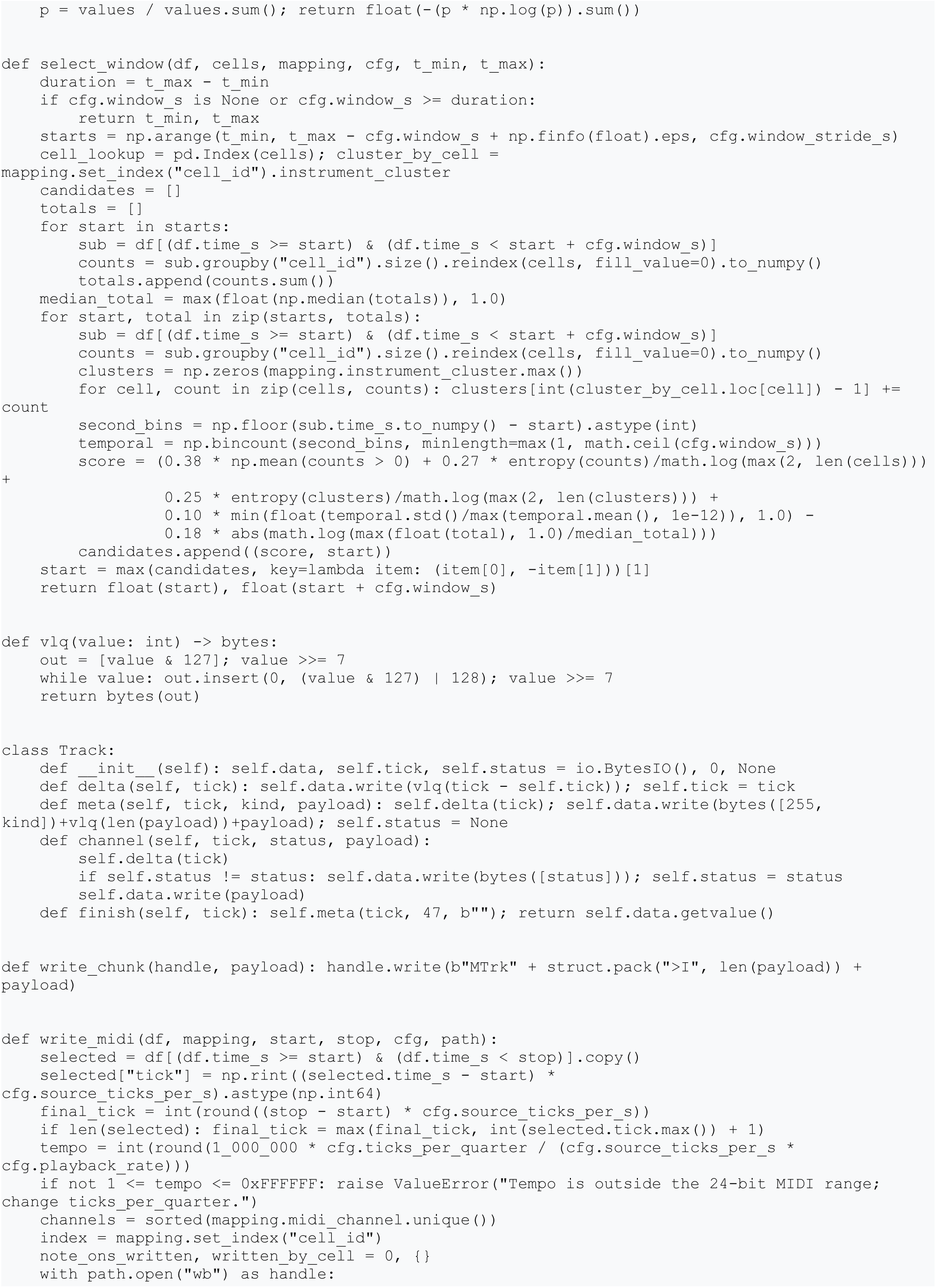
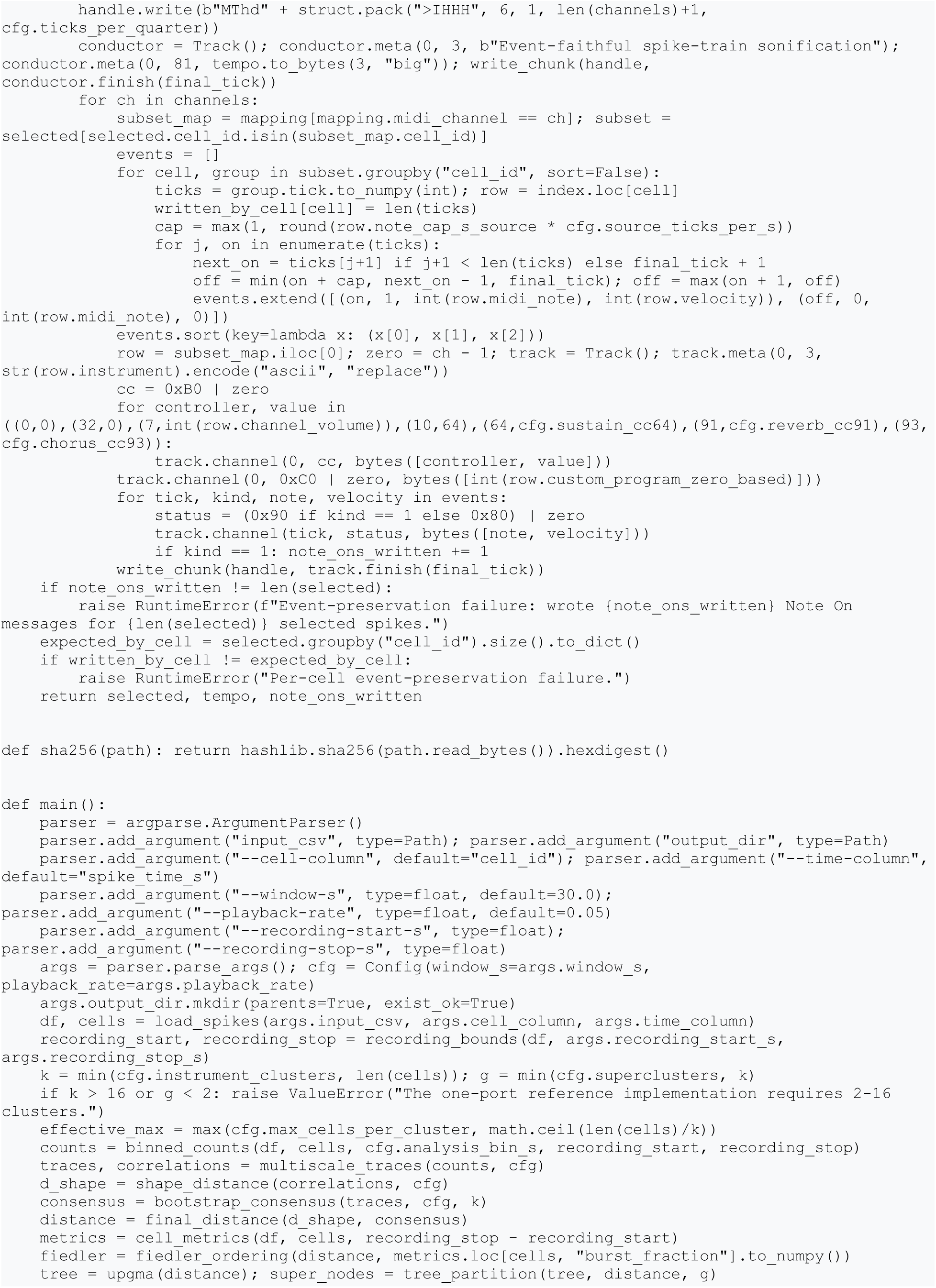
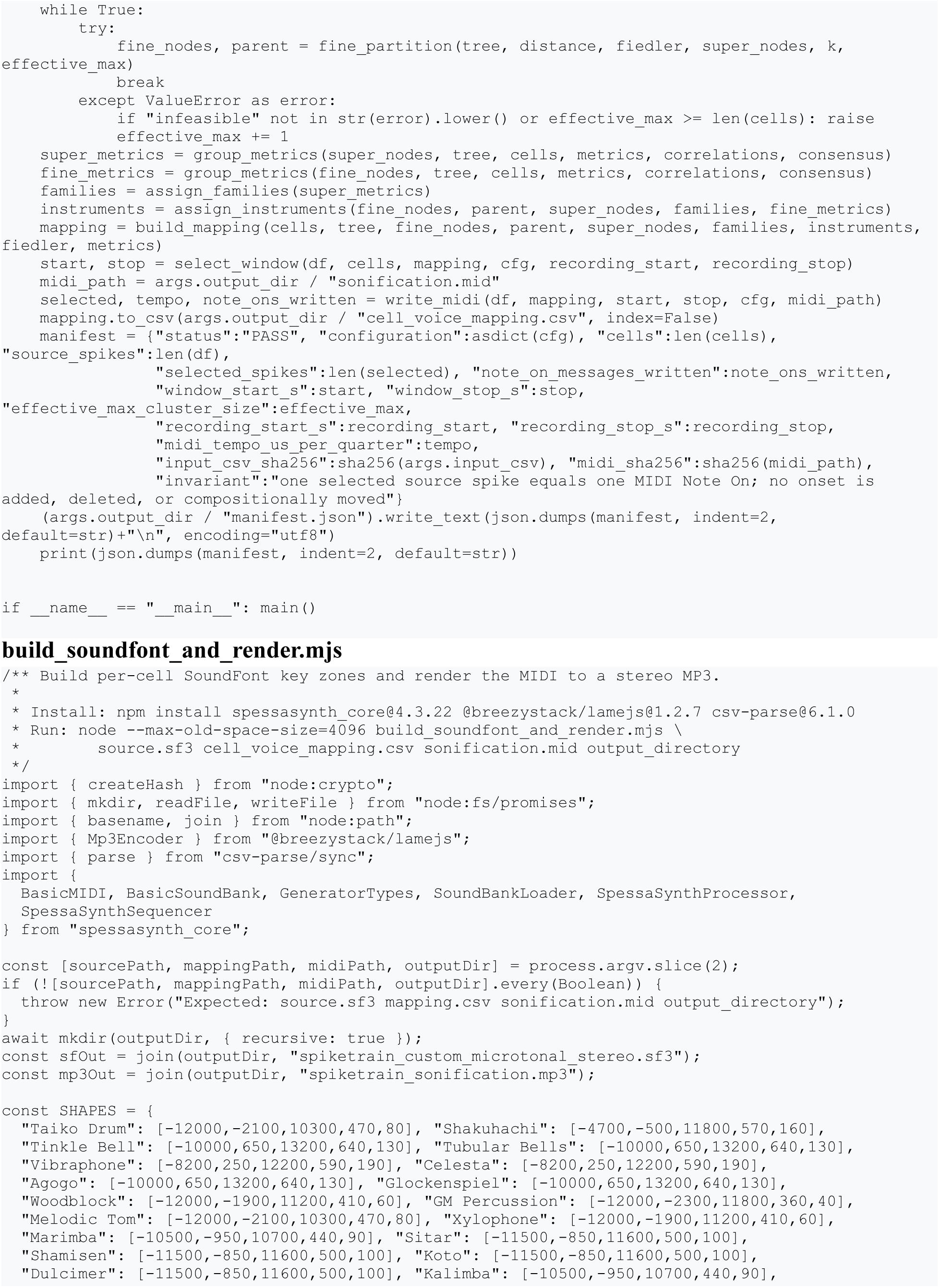
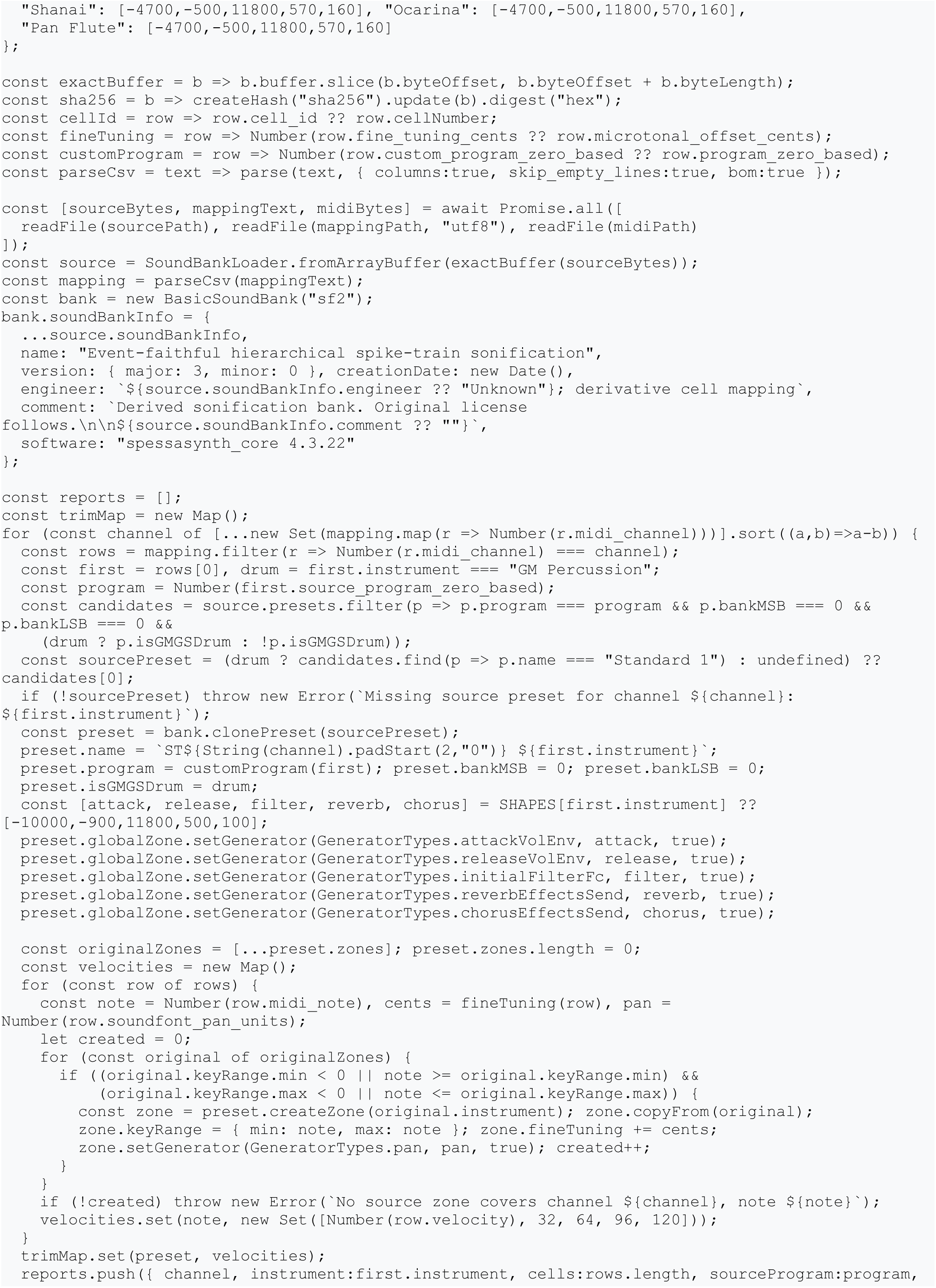
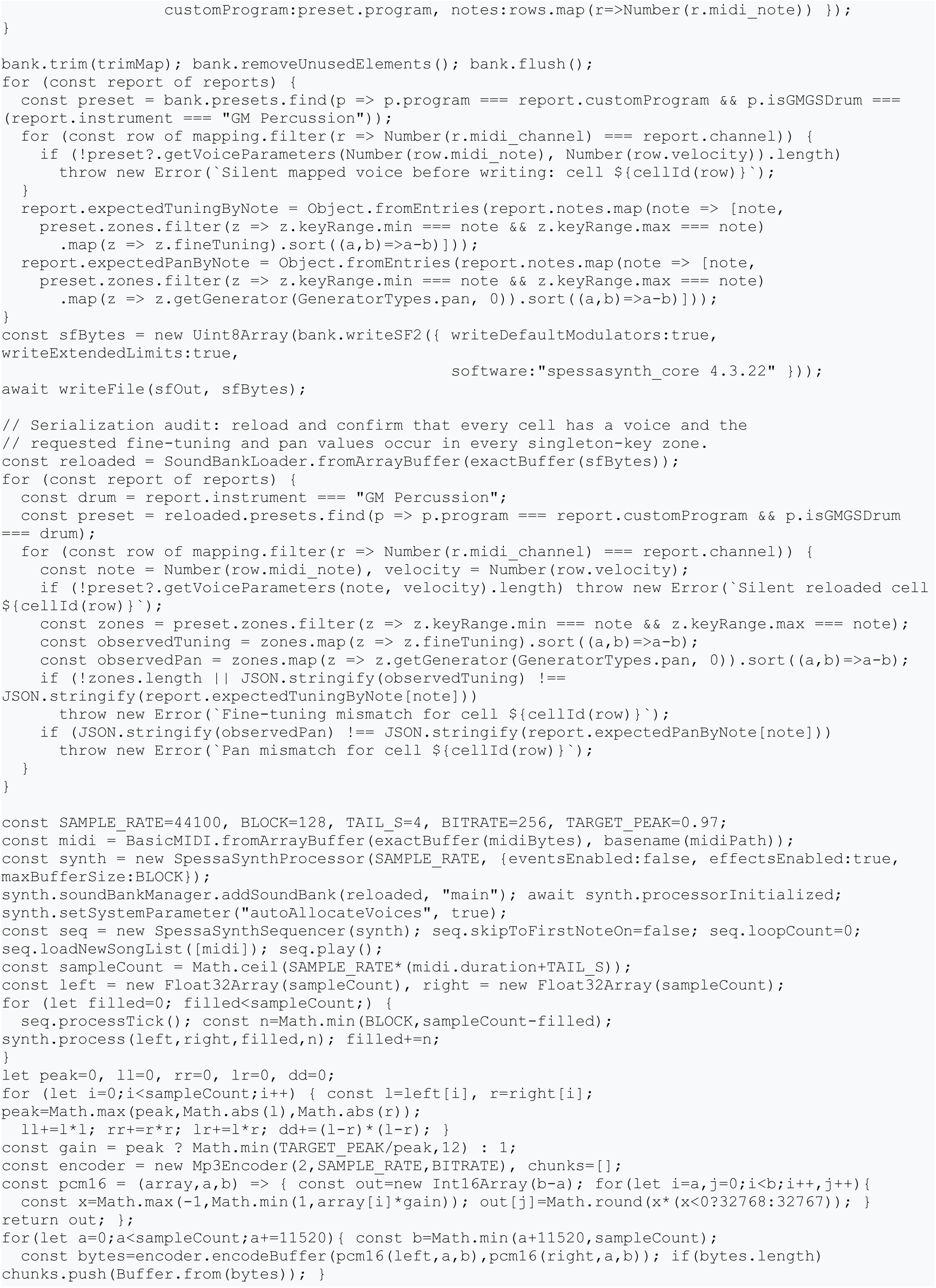
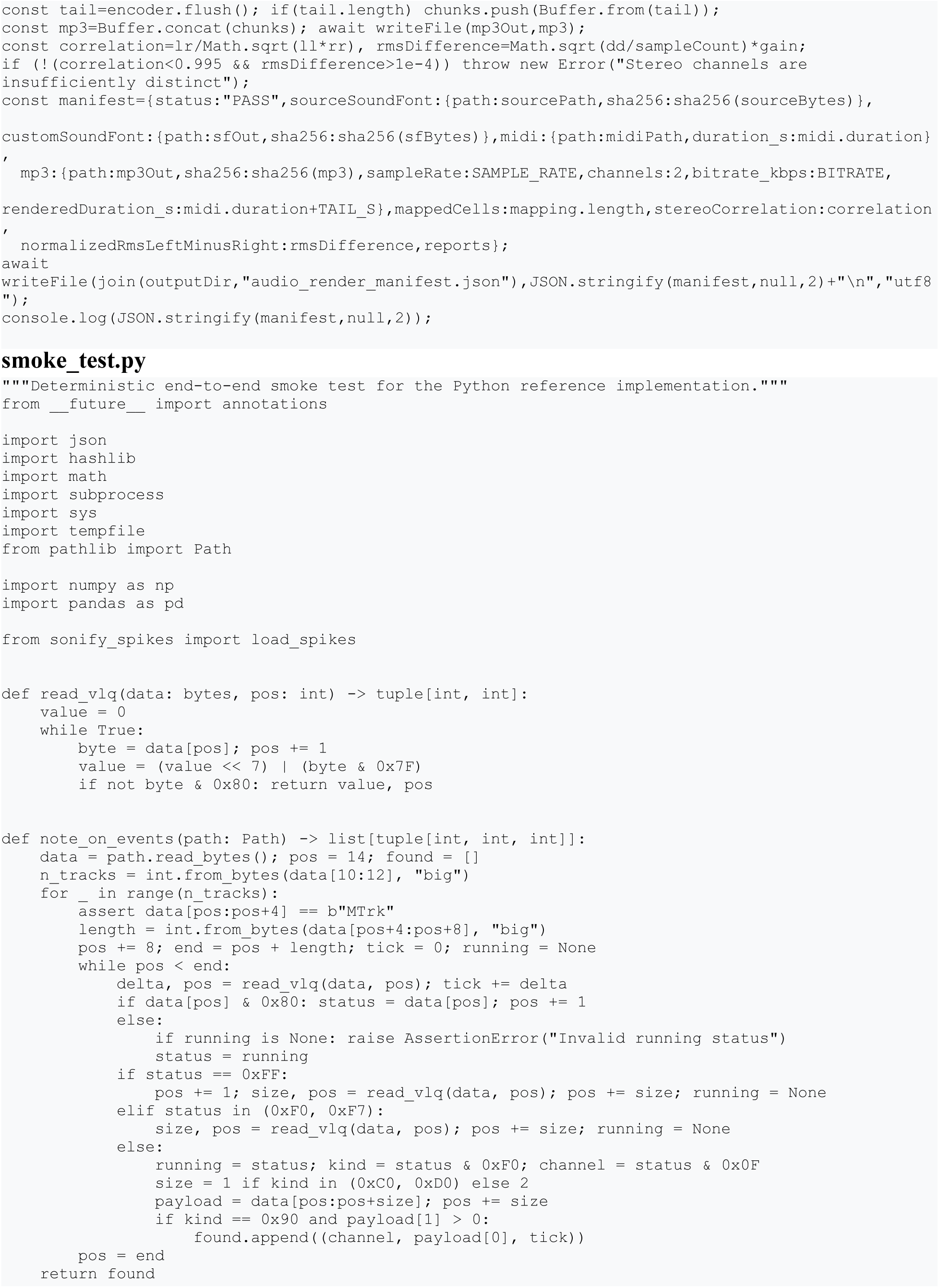
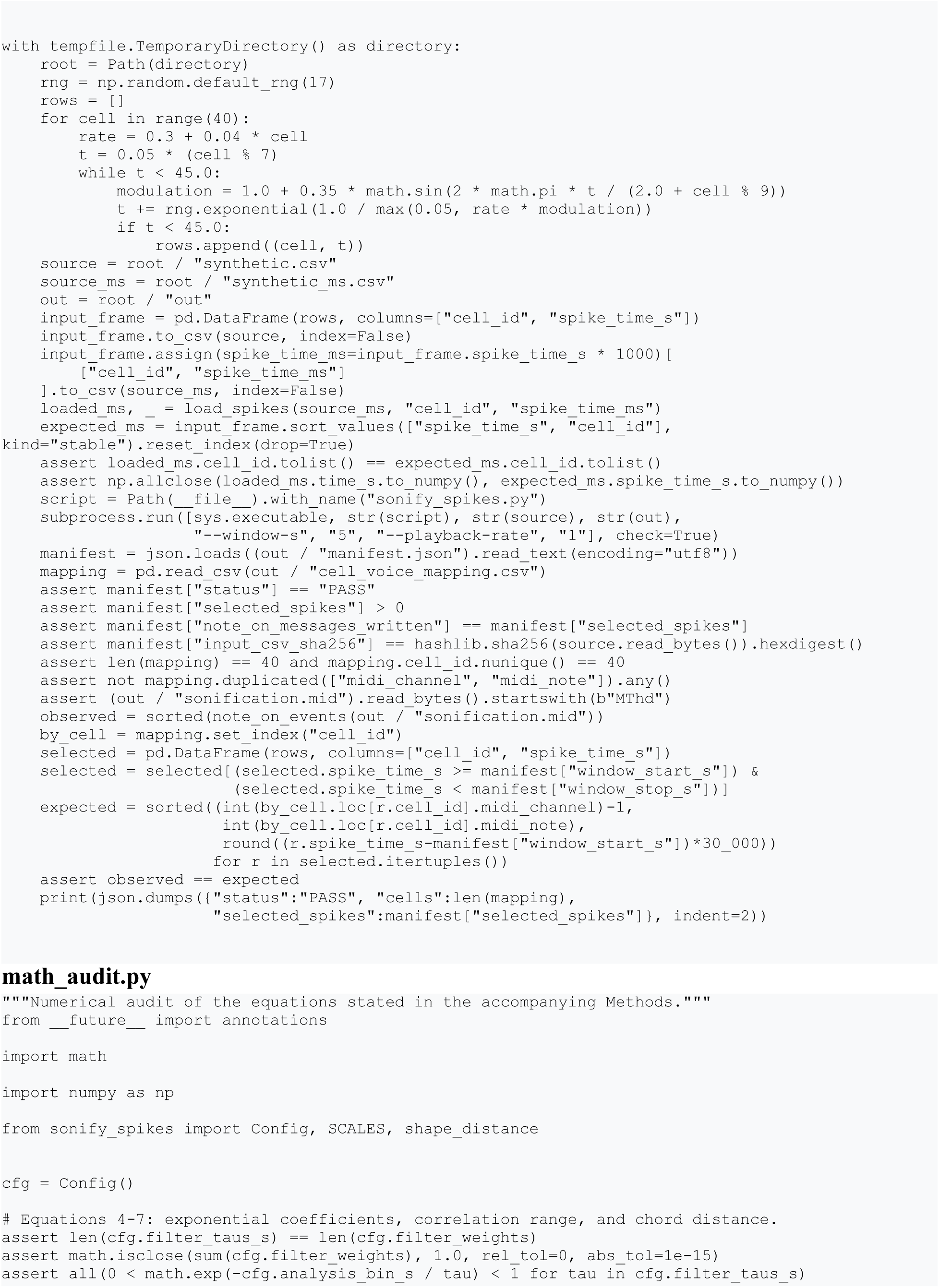
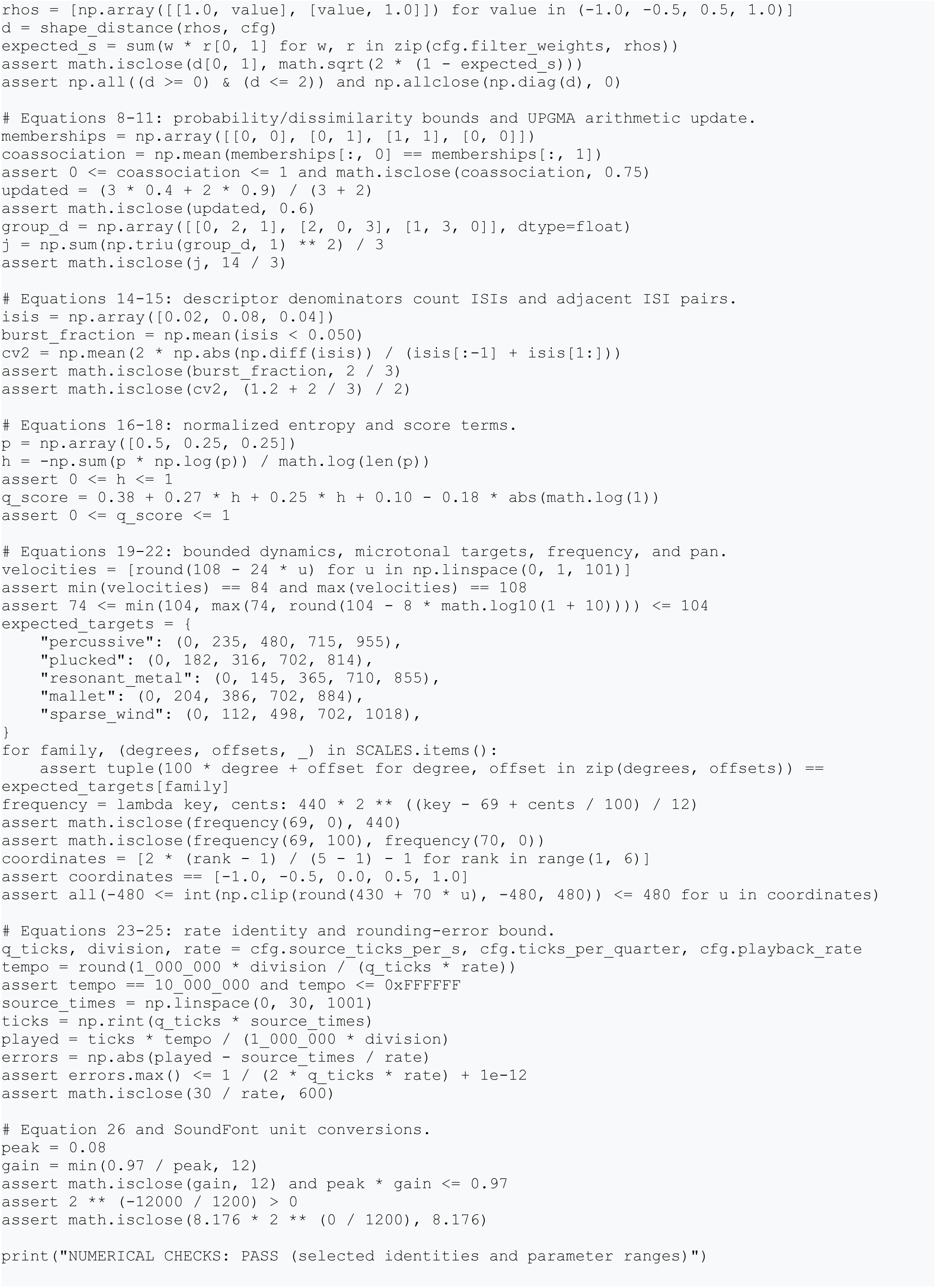

## Notes

https://github.com/etangval/HERO

https://www.youtube.com/watch?v=zOYGXcp727A

### Competing Interest Statement

The authors have declared no competing interest.

